# GLOWORM-META: Modelling gastrointestinal nematode metapopulation dynamics to inform cattle biosecurity research

**DOI:** 10.1101/2025.02.21.639461

**Authors:** Olivia K. Ingle, Lynsey A. Melville, Sam Beechener, Claire Hardy, Alison Howell, Neil P. Hobbs, Niamh Mahon, Eric R. Morgan, David Bartley, Hannah Rose Vineer

**Affiliations:** University of Liverpool, Department of Infection Biology & Microbiomes, Institute of Infection, Veterinary and Ecological Sciences, Leahurst Main Building, Neston, Cheshire, CH64 7TE, England; Moredun Research Institute, Pentlands Science Park, Bush Loan, Penicuik, EH26 0PZ, Scotland; Scotland’s Rural College, Peter Wilson Building, The King’s Buildings, West Mains Road, Edinburgh, EH9 3JG, Scotland; The James Hutton Institute, Craigiebuckler, Aberdeen, AB15 8QH, Scotland; Queen’s University Belfast, School of Biological Sciences, 19 Chlorine Gardens, Belfast, BT9 5DL, Northern Ireland

**Keywords:** Gastrointestinal nematodes, Metapopulations, Cattle, Livestock management, Parasite

## Abstract

Gastrointestinal nematode (GIN) parasite infections in grazing livestock cause significant disease, and are responsible for estimated annual losses of over €1.8 billion in Europe alone. The management of GINs in cattle is threatened by anthelmintic drug resistance (AR). Immediate action is needed to slow the progression of AR in cattle GINs and avoid the increasingly common scenario of multiple drug resistance seen in sheep. Although AR can arise independently on multiple farms, it may also be spread between holdings via purchased cattle. Therefore, effective biosecurity measures on cattle enterprises could help to reduce the risk of establishment of AR populations. A metapopulation model was developed and validated for two GIN species infecting cattle, *Ostertagia ostertagi* and *Cooperia oncophora*, incorporating the full parasite life cycle, weather- and immunity-dependent parasite life history traits and multiple pasture sub-populations. This allowed for complex grazing management strategies and weather influences to be simulated. The models successfully replicated the seasonal patterns and intensity of infections reported in multiple published longitudinal datasets. Global sensitivity analysis against four Quantities of Interest (QoIs) related to factors affecting the safety of the resident herd and of the purchased animals was used to quantify the influence of candidate biosecurity measures. The duration of quarantine, the date of purchase (weather/seasonal influences) and the intensity of infection on the day of purchase strongly influenced the QoIs. The outcomes for the UK were not significantly influenced by the geographic location of the purchasing farm, suggesting that the influence of weather patterns on GIN populations outweighs that of regional climate differences, and thus regional variations to GIN biosecurity recommendations are not warranted without alternative evidence to support this. The model presented here is the first full lifecycle GIN metapopulation model for *O. ostertagi* and *C. oncophora*, validated against longitudinal field data, and can be broadly used to evaluate the relative efficacy of a range of cattle GIN management strategies, as demonstrated here. These findings offer valuable insights to focus initial biosecurity recommendations for cattle enterprises, and are being used to direct qualitative and quantitative research to refine recommendations.

**Highlights:**

- A metapopulation model for cattle GINs was developed
- The model includes weather-dependence and complex grazing
- The model replicated seasonal infection patterns from multiple datasets
- Sensitivity analysis explored influential biosecurity measures
- The influence of weather outweighed that of geographic location (climate)
- Initial recommendations for biosecurity and further research are made

## 1. Introduction

Gastrointestinal nematodes (GINs) are a major threat to the health and productivity of cattle, resulting in significant economic burdens amounting to estimated annual losses of over €1.8 billion in Europe alone (Charlier et al., 2020). *Ostertagia ostertagi* and *Cooperia oncophora* are the predominant GINs for cattle residing in temperate regions. Their management is threatened by anthelmintic drug resistance (AR; (Papadopoulos et al., 2012; Kaplan, 2020; Rose Vineer et al., 2020a). Immediate action is needed to slow the progression of AR in cattle GINs and avoid the increasingly common scenario of multiple drug resistance seen in the sheep industry (Jackson et al., 2011; Martínez-Valladares et al., 2015). AR can independently arise on multiple farms, due to repeated use of anthelmintics leading to selection pressures which allow resistant worms to survive and reproduce. This process is often exacerbated by under-dosing, frequent treatments, and inappropriate rotation between anthelmintic classes (Smith et al., 1999). In this case, considered use of anthelmintics is required to preserve efficacy (Bates et al., 2012). However, AR may also spread between holdings via purchased cattle. Research to quantify the risk of spread between cattle farms is ongoing, while the World Organisation for Animal Health recommend quarantine with anthelmintic treatment and confirmation of efficacy by faecal egg count (FEC) to prevent the introduction of resistant GINs (for Animal Health, 2019). This is in line with COWS (Control of Worms Sustainably) recommendations for general GIN biosecurity in the UK (COWS, 2021). Thus, biosecurity measures on cattle enterprises aiming to reduce the population of GINs introduced onto a farm are also appropriate for reducing the risk of establishment of AR populations, provided the anthelmintics used at this time are effective. However, the WOAH and COWS recommendations currently lack specificity, and the the evidence base required to provide more detailed guidance on biosecurity factors such as the duration and location of quarantine (acknowledging the common impracticalities of lengthy periods of housing/yarding of bought-in stock), and interactions with local environmental conditions, is lacking.

Epidemiological models can guide the design of effective and sustainable control strategies (Szmaragd et al., 2009; Mayo et al., 2020; Lubinda et al., 2021), and are increasingly used to support the design of sustainable livestock parasite control recommendations (Rose et al., 2015; Rose Vineer et al., 2020b). The population dynamics of GINs arise from complex host-environment-parasite interactions (Wang et al., 2022), and epidemiological models provide the tools to explore such interacting factors with minimal resources and risk (Vineer, 2020), in a way that can be generalised across farm contexts. A range of cattle nematode models has been developed (reviewed by Verschave et al., 2016) Over the previous decade, two GLOWORM frameworks representing the free-living and parasitic stages of trichostrongylid nematodes respectively, have been developed and extended to model the population dynamics of a range of nematode species infecting sheep and cattle (Rose et al., 2015; Rose Vineer et al., 2020b; McCarthy et al., 2022). These models were developed to simulate the impact of GLObal change on parasitic WORMs, especially the impact of weather patterns and anthropogenic interventions (such as veterinary treatments and grazing management) on the population dynamics of ruminant GINs. The model of the free-living stages, GLOWORM-FL, tracks the development, mortality and migration of the eggs and free-living larvae of GINs on pasture (Rose et al., 2015). The parasitic stages model, GLOWORM-PARA, tracks the infection of ruminant hosts by the parasitic stages of GINs, and the subsequent development, mortality, and egg production of the GINs (Rose Vineer et al., 2020b). The models take advantage of technological advances (e.g. Wang et al., 2020) and the increasing availability of longitudinal data for validation, and have been successfully applied to a range of systems to inform GIN management such as rotational grazing and contamination mapping (McFarland et al., 2022), and relative contributions of host species to shared GIN populations in multi-host systems (Khanyari et al., 2024; Walker et al., 2018). However, as they each only simulate half of the GIN life cycle, they require detailed longitudinal parasitological data as input, which limits their applications. To model a wider range of parasite transmission patterns and evaluate the relative impact of a range of candidate control measures, a full life-cycle model of parasite epidemiology is required. Furthermore, GLOWORM-FL has been previously parameterised to simulate only *Ostertagia ostertagi* dynamics in cattle (plus *Haemonchus contortus* and *Teladorsagia circumcincta* in sheep), while GLOWORM-PARA has been previously parameterised and validated for both cattle GIN species. Thus, parameters also need to be defined for *C. oncophora* free-living stages.

Therefore, the aim of the work presented here was to identify the most impactful biosecurity practices for GIN control in cattle, by developing a validated, full-life cycle model for *O. ostertagi* and *C. oncophora*, incorporating herd management and anthelmintic treatment. A metapopulation model, GLOWORM-META, was developed to combine the existing frameworks described above for the free-living GIN stages (GLOWORM-FL, Rose et al. (2015)) and the parasitic within-host stages (GLOWORM-PARA, Rose Vineer et al. (2020b)), and track GIN sub-populations on pastures coupled by herd rotations. Global sensitivity analysis was used to evaluate the interacting effects of climate, weather and a range of biosecurity measures, to focus further research.

## 2. Methods

### 2.1. Modelling GIN population dynamics – the GLOWORM-META framework

#### 2.1.1. Model description

The GLOWORM-META framework was developed to simulate the population dynamics of *O. ostertagi* and *C. oncophora* infecting cattle that are rotationally grazed, by combining and extending the GLOWORM-FL (Rose et al., 2015) and GLOWORM-PARA (Rose Vineer et al., 2020b) model frameworks (Figure 1; GLOWORMFL and GLOWORM-PARA are referred to hereafter as modules to avoid confusion with the full life-cycle model, GLOWORM-META). To develop the full lifecycle framework, the modules were linked by host ingestion of third-stage GIN larvae on herbage, and by deposition of GIN eggs onto pasture in faeces (Figure 1).

**Figure 1:**
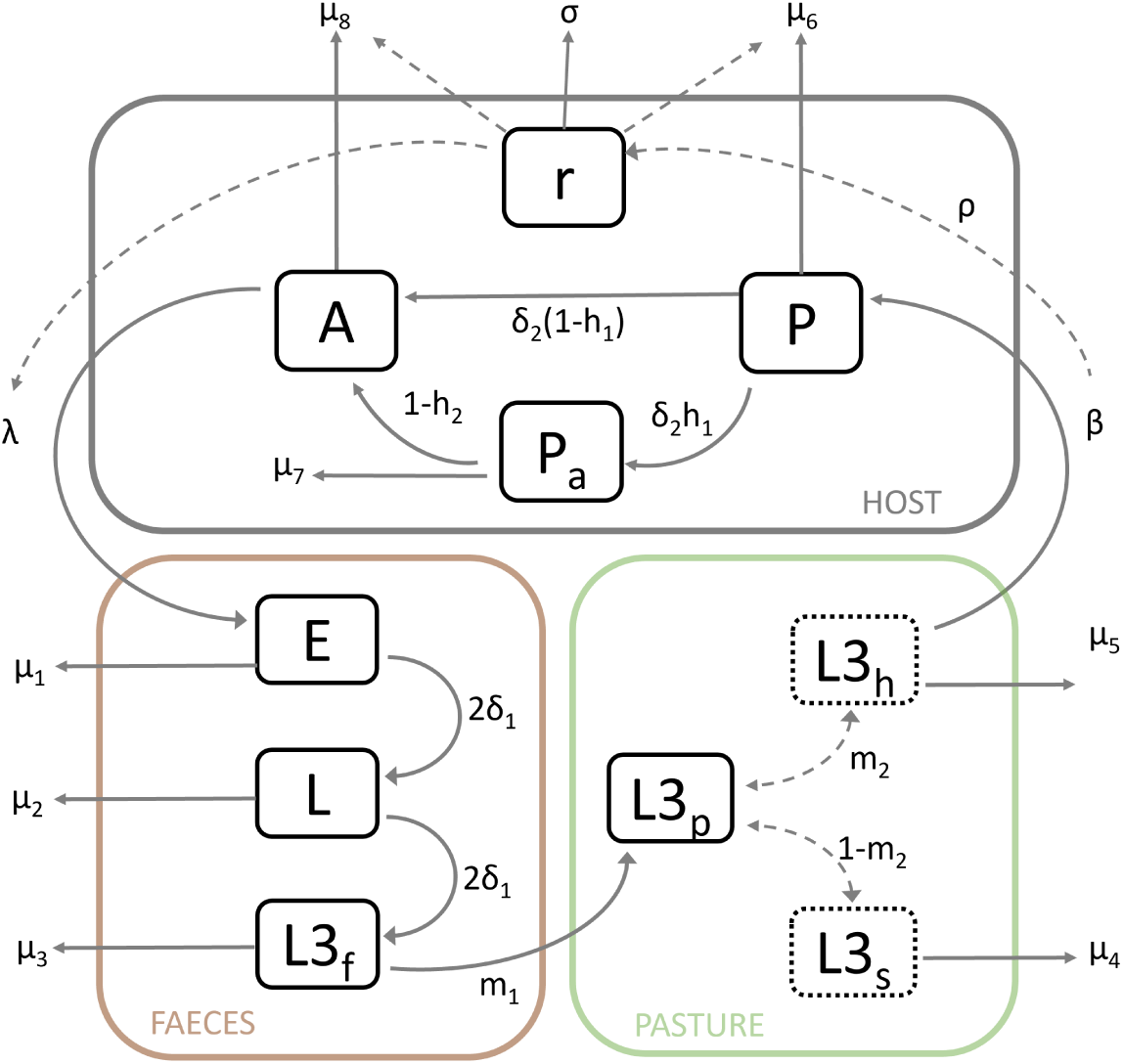
The conceptual framework of the full life cycle model which forms the basis of the GLOWORM-META model. This model combines the GLOWORM-FL module (Rose et al., 2015; faeces (brown) and pasture (green) components) with the GLOWORM-PARA module (Rose Vineer et al., 2020; host component (grey)), linked by host ingestion of third-stage larvae (*β*) and egg excretion (*λ*). Model parameters (shown in boxes) and state variables are described in Table 1. Briefly, Eggs (*E*) are deposited on pasture in faeces and develop to third stage larvae in faeces (*L*3*_f_*), which then migrate horizontally onto pasture and migrate vertically between the herbage (*L*3*_h_*) and soil (*L*3*_s_*). L3 on herbage (*L*3*_h_*) are ingested by the host and exsheath to join the pool of pre-adult parasitic stages (*P*). These stages either develop to adults (*A*) directly, or arrest their development (*Pa*; hypobiosis). Host immunity (*r*) is dependent on exposure to infection, and influences the mortality and fecundity of the parasitic stages. (*L*3*h*) and (*L*3*s*) are shown using dotted boxes because they are subcomponents of the total L3 on pasture (*L*3*p*). Transitions are shown as solid arrows, while dependencies between state variables and parameters are shown as dotted arrows.

**Table 1:** Definitions of state variables, input variables and parameters, and assigned values/functions for *O. ostertagi* and *C. oncophora* infecting cattle. All rates are instantaneous daily mortality rates, unless otherwise stated.

| Symbol | Description | *O. ostertagi* | *C. oncophora* |
| --- | --- | --- | --- |
| $E$ | Eggs in faeces | Equation 1 | |
| $L$ | L1 and L2 in faeces | Equation 2 | |
| $L3_f$ | L3 in faeces | Equation 3 | |
| $L3_p$ | L3 on pasture | Equation 4 | |
| $P$ | Pre-adult parasitic stages | Equation 5 | |
| $Pa$ | Hypobiotic larvae | Equation 6 | |
| $A$ | Adults | Equation 7 | |
| $r$ | Host resistance (immunity) | Equation 8 | |
| $T$ | Daily mean air temperature ($^{\circ}C$) | Input | |
| $R$ | Daily total rainfall (mm) | Input | |
| $c$ | Daily kg dry matter intake | Input | |
| $B$ | Standing biomass, $kgDMha^{-1}$ | Input | |
| $H$ | Host stocking density $ha^{-1}$ | Input | |
| $D$ | Hours of daylight | estimated using input coordinates and *geosphere* R package | |
| $ET$ | Daily potential evapotranspiration (mm) | $0.55D \times \frac{4.95e^{0.062T}}{100}$ | |
| $\beta$ | Host herbage ingestion rate | $-\ln(1 - \frac{c}{B})$ | |

| Continuation of Table 1 |  |  |  |
| --- | --- | --- | --- |
| Symbol | Description | *O. ostertagi* | *C. oncophora* |
| $d$ | Driver of hypobiosis | $\delta_1$ | |
| $\delta_1$ | $E$ to $L3_f$ development rate | $-0.07258 + 0.00976T$ | $-0.0401 + 0.00821T$ |
| $\delta_2$ | $P$ to $A$ development rate | 0.041 | 0.041 |
| $\mu_1$ | $E$ mortality rate | $e^{-4.38278-0.1064T+0.0054T^2}$ | $e^{-4.38278-0.1064T+0.0054T^2}$ |
| $\mu_2$ | $L$ mortality rate | $e^{-4.38278-0.10640T+0.00540T^2}$ | $e^{-4.38278-0.10640T+0.00540T^2}$ |
| $\mu_3$ | $L3_f$ mortality rate | $e^{-4.864-0.206T+0.0048T^2+0.00008T^3}$ | $e^{-4.726-0.204T+0.0044T^2+0.00009T^3}$ |
| $\mu_4$ | $L3_s$ mortality rate | $e^{-5.743-0.1755T+0.0068T^2-0.00003T^3}$ | $e^{-5.822-0.1749T+0.007T^2+0.00002T^3}$ |
| $\mu_5$ | $L3_h$ mortality rate | $\mu_3$ | |
| $\mu_6$ | $P$ mortality rate | $0.054 + (0.086 - 0.054)r$ | $0.044 + (0.12 - 0.044)r$ |
| $\mu_7$ | $Pa$ mortality rate | 0.002 | |
| $\mu_8$ | $A$ mortality rate | $0.028 + (0.08 - 0.028)r$ | $0.04 + (0.12 - 0.04)r$ |
| $m_1$ | $L3_f$ horizontal migration rate | $\begin{cases} e^{-3.296+0.046R}, & R \geq 5 \\ 0, & R < 5 \end{cases}$ | $\begin{cases} e^{-3.35+0.056R}, & R \geq 6 \\ 0, & R < 6 \end{cases}$ |
| $m_2$ | $L3_p$ vertical migration rate | $(e^{-5.482+0.454T-0.013T^2}) \times 0.27$ | $(e^{-5.482+0.454T-0.013T^2}) \times 0.46$ |
| $\lambda$ | Per $A$ fecundity rate | $4.96 - (4.96 - 4.58)r$ | $7.30 - (7.30 - 6.44)r$ |
| $h_1$ | Hypobiosis rate | $0.3 - ((0.3 - 0.02)/d_{\max})d_t$ | $0.06 - ((0.06 - 0)/d_{\max})d_t$ |
| $h_2$ | Emergence rate | $(1/(d_{\max} - d_{\min})) \times (d_t - d_{\min})$ | |
| Symbol | Description | *O. ostertagi* | *C. oncophora* |
| $\rho$ | Immune response rate per *L3* ingested | $5.98e^{-5}$ | $1.3e^{-4}$ |
| $\sigma$ | Immune decay rate | 0.002 | |
| $C_e$ | Egg correction factor | 1 | |

The resulting full life-cycle model was then extended to incorporate spatially implicit metapopulation dynamics representing sub-populations of free-living stages on different pastures (also referred to as patches) coupled by the movements of a single group of hosts between the pastures. The free-living stages module (green and brown in Figure 1) independently simulates the population dynamics of the sub-populations on each of six pastures (patches), which are represented by *i*, with *i* = 1, 2, 3, 4, 5, 6. All hosts move as a single herd between the available pastures (the metapopulation network). The location of the herd, *j*, with *j* = 0, 1, 2, 3, 4, 5, 6, determines which patch is coupled with the parasitic stages module (grey in Figure 1). Because all hosts move between patches in unison, state variables and parameters denoted by *ij* in the equations below represent either decoupling of the free-living stages on patch *i* and the parasitic stages in the host when *i ̸*= *j*, or coupling when *i* = *j*. In the latter case, parameters and state variables are as defined in Table 1 and as simulated, respectively, allowing eggs, *E*, to be deposited onto a pasture (Eq. 1) and infective larvae on herbage, *L*3*_h_*, to be ingested from a pasture (Eq. 4, 5 and 8). When *i ̸*= *j*, parameters and state variables with the subscript *ij* in the equations below are equal to zero. Where *j* = 0, this represents complete uncoupling, for example when animals are housed or kept on hard standing. State variables and parameters are summarised in Table 1.

#### 2.1.2. Model equations

GIN eggs and larvae develop and die on pasture at stage- and environment-specific temperature-dependent rates (Table 1; *δ*_1_ and *µ*_1-5_). New eggs (*E*) are deposited on pasture based on the daily fecundity rate of the GIN species (*λ*), the number of Adult GINs per host (*A*), and host stocking rate per hectare (*H*). Eggs excreted can also be subject to moisture deficit-dependent mortality (*C_e_*; Equation 1), although this is not used in the present model due to the high water content of cattle dung and the rapid formation of a protective crust on the surface (Rose, 1961). For *C_e_*, and also to derive moisture-dependent horizontal migration parameter (*m*_1_; Table 1; Rose et al., 2015), potential evapotranspiration (*ET*) is estimated using the Hamon method (Xu and Singh (2001); Table II). The development rate is doubled as it is a daily rate applied across two life-cycle stage transitions (Equations 1-3).

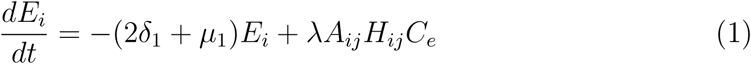

Larvae develop through first and second pre-infective larval stages (*L*; Equation 2) to third-stage infective larvae in faeces (*L*3*_f_*; Equation 3).

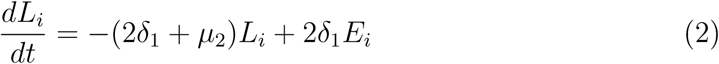

*L*3*_f_* migrate horizontally out of faeces at rate *m*_1_ based on temperature and rainfall. They then migrate vertically between herbage and soil based on a temperature-dependent vertical migration rate (*m*_2_), where they are subject to different mortality rates.

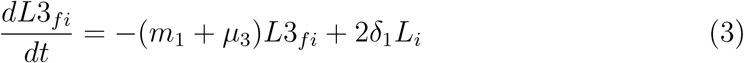

A proportion of *L*3*_p_* migrate onto herbage (*m*_1_) and are removed by the grazing host. The rate of removal is dependent on the daily rate of herbage ingestion per host (*β*), which is determined by the daily dry matter intake (*c*) as a proportion of the standing biomass per hectare (*B*; Rose et al., 2016), which can be used to account for the impact of interactions between GIN vertical migration behaviour and cattle grazing heights on the proportion of L3 on herbage that are ingested, or supplementary feeding (*C_g_*; Table 1).

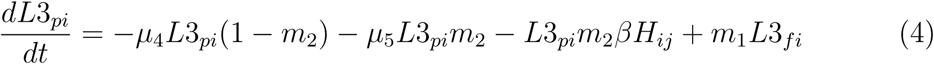

The ingested L3 then enter the host, where they are represented by the pre-adult parasitic (*P*) state variable (Eq. 5). Here, they either develop (*δ*_2_) to the adult stage (*A*; Eq. 7) or enter hypobiosis (*Pa*; Eq. 6) at a rate determined by the hypobiosis parameter (*h*_1_). Hypobiotic larvae resume development at a rate determined by the emergence parameter (*h*_2_).

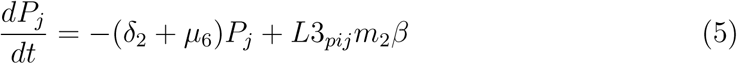

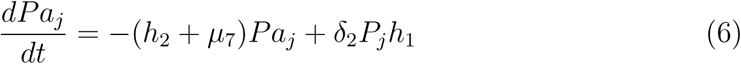

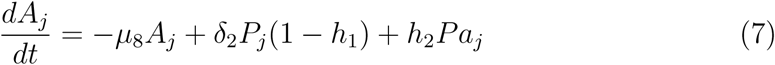

Host resistance to infection (immunity; *r*), increases with exposure to ingested L3 (*ρ*), and wanes at a constant rate (*σ*).

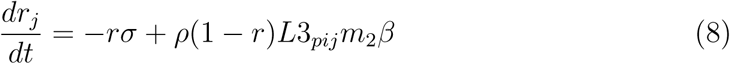

The numbers of *L*3*_p_* residing on herbage (*L*3*_h_*) and in soil (*L*3*_s_*) are not explicitly simulated as state variables, but are included in equations 4, 5 and 8, and can be calculated from the daily output of simulations using the equations:

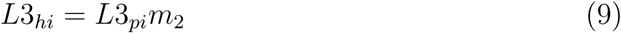

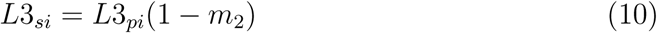

#### 2.1.3. Environmental input data and model implementation

The model detailed above is implemented in R (v 4.3.0 R Core Team (2023)) using the *deSolve* package (Soetaert et al., 2010) as described for the modules (Rose et al., 2015; Rose Vineer et al., 2020).

Daily rainfall (mm) and air temperature (*^◦^*C) data are required, corresponding to the simulation length. Start and end dates, and latitude and longitude data are required to obtain daylength data (hours) using the *daylength* function of the *geosphere* package (Hijmans, 2024). The present framework assumes that all patches are nearby and that a single weather dataset adequately represents the ambient conditions at all of the patches. The weather data are used to estimate the temperature- and rainfall-dependent variables (Table 1) which are introduced to the model as seasonally forced variables.

Equations 1-4 above, representing the free-living stages, are replicated for each patch (*i*). The location of the host group (*j*) is input at the start of the simulations with corresponding dates representing the movements between pastures (patches). The location data are then used to generate a binary variable for each patch indicating whether the host group is grazing that patch (1) or not (0). Daily stocking rates, and a single estimate of herbage biomass are also required.

Throughout model development, the programmers practised continuous integration and informal unit testing of model code as complexity increased. Formal unit testing was conducted on the final model to further protect against programming errors introduced with the additional complexity of the linked metapopulations. For this, the model was run for Herd A in the McFarland dataset detailed under “Model validation” below, without pasture rotations. This was repeated for each patch in the model and output tested for differences using the *identical* function of the *testthat* R package (Wickham, 2011). Following peer-reviewed publication, all code will be available at - https://github.com/hannahvineer/gloworm-meta.

### 2.2. GLOWORM parameters

All parameters are defined in Table 1.

#### 2.2.1. Free-living stages

The daily temperature-dependent development rate for *C. oncophora* is as described by Sauermann and Leathwick (2018). Mortality rates for the free-living stages of *C. oncophora* are as described in Wang et al. (2022). All development and mortality rates for the free-living stages of *O. ostertagi* are as described in Rose et al. (2015), except the mortality of L3 in soil and on herbage, which were taken from Wang et al. (2022).

Instantaneous daily vertical migration rates for *C. oncophora* and *O. ostertagi* at a range of rainfall volumes (mm) were extracted from Wang et al. (2022) using PlotDigitizer (https://plotdigitizer.com/). A log-linear model was fit to these rates to provide an equation that could be used to estimate horizontal migration rates from rainfall (*O. ostertagi*; a = −3.230, b = 0.0460, *R*^2^ = 0.574, *R*^2^*adj* = 0.553, *F*_1,20_ = 26.9, p *<* 0.001, *C. oncophora*; a = −3.35, b = 0.0556, *R*^2^ = 0.588, *R*^2^*adj* = 0.566, *F*_1,19_ = 27.1, p *<* 0.001). We used this approach in lieu of the equations provided in Wang et al. (2022) as the linear model fit in Wang et al. (2022) estimated zero migration at rainfall values *<*5mm, which is not realistic for simulations in humid temperate environments such as the UK, where cattle dung is likely to dry at a slower rate than in Alberta, Canada, where Wang et al. made their observations.

To estimate the vertical migration rates it was necessary to conduct additional *in vitro* observations. Faeces containing *O. ostertagi* or *C. oncophora* eggs were obtained from calves infected with either *O. ostertagi* (MOo2; isolated in 1995 from Belgium and passaged through parasite free cattle annually) or *C. oncophora* (MFie26; isolated in 2021 from a field sample), respectively. Larvae were cultured to L3 at 15°C for 10 days, and extracted from faeces using a modified Baermann method. Small pots (3.5cm diameter, 4.5cm depth) containing 4.5cm of soil and *Lolium perenne* grass 5 cm in height were seeded with approximately 500 larvae in water, pipetted onto the soil surface, grass pots were moistened with 1ml of tap water sprayed onto the leaves and incubated at a range of temperatures between 5-25°C (8 replicates per treatment). After 24 hours, larvae were recovered from half of the replicates, and after 48 hours the same process was followed for the other half. Grass leaves were cut at soil level using sterilized scissors and soaked overnight at ambient room temperature in small individual pots containing tepid tap water and a drop of Tween-20. Leaves were removed from the pots and the liquid was settled for 2 hours before the volume was reduced to 5ml. The remaining liquid was washed into a 12ml polyallomer tube (Beckman Coulter Inc, USA), centrifuged at 200*×*g for 2 minutes and the supernatant removed. Saturated potassium iodide (10ml) was added and tubes were centrifuged again. As larvae float in saturated potassium iodide, tubes were clipped below the meniscus using artery forceps to isolate larvae and the contents of the top chamber was washed into a cuvette for analysis and larval counting under *×*40 magnification. After correcting for estimated recovery rates (Wang et al., 2022), the proportions of larvae migrating onto herbage ranged between 0-0.23 for *O. ostertagi* and 0-0.41 for *C. oncophora*. Pairwise Kolmogorov-Smirnoff tests (Doe et al., 2024), confirmed that, on the whole, the distribution of the data did not differ between temperature treatments, with the exception of the 15*^◦^*C treatment differing significantly from the 20*^◦^*C (p = 0.007) and 25*^◦^*C (p = 0.001) treatments for *C. oncophora*, and the 10*^◦^*C treatment differing significantly from the 25*^◦^*C (p = 0.022) treatment for *O. ostertagi*. Therefore, it was not appropriate to fit a linear model to these data to estimate temperature-dependent migration rates. However, the data do indicate that migration rates are higher at 20-25*^◦^*C, consistent with Rose et al.’s (2015) model of sheep nematode vertical migration, and it is possible that the small numbers of nematodes recovered (ranging from 0-41 per replicate) and the measurement error associated with nematode recovery from herbage (Lancaster, 1970), are confounding any patterns in the present dataset. Therefore, the vertical migration model defined by Rose et al., (2015) for sheep nematodes was calibrated for *O. ostertagi* and *C. oncophora* using the *optim* function in R to apply a scaling factor which minimises the sum of squared errors between the empirical data and the vertical migration model equation. This resulted in a scaling factor of 0.092 for *C. oncophora* and 0.054 for *O. ostertagi* (Table 1).

#### 2.2.2. Parasitic stages

Parameters for the parasitic stages and immune response, except the rates described below, are as described in Rose Vineer et al. (2020) for both *O. ostertagi* and *C. oncophora*.

Instantaneous daily minimum and maximum Pre-adult mortality rates for *O. ostertagi* were estimated using equation 5, the mean and minimum establishment rates reported in Verschave et al. (2014), respectively, and a prepatent period of 17 days (Rose Vineer et al., 2020). Maximum instantaneous daily adult mortality rates for *C. oncophora* were approximated from the upper limits of data collated by meta-analysis ((Verschave et al., 2014, 2016; Verschave, 2015)

Minimum and maximum proportions of ingested L3 entering hypobiosis were determined for both species from the minimum and approximate maximum arrest rates in published meta-analyses (Verschave et al., 2014, 2016; Verschave, 2015).

### 2.3. Model validation

The free-living (GLOWORM-FL) and parasitic (GLOWORM-PARA) modules that were combined in this model have been previously validated as independent units. The free-living module was validated using data from Canada and the UK (Rose et al., 2015; Rose Vineer et al., 2020b; Wang et al., 2022), while the parasitic stages module was validated for set-stocked calves in their first grazing season in Belgium (Rose et al., 2015)

Validation of the complete life-cycle model presented here, incorporating rotational grazing (GLOWORM-META) and the updated parameter set, was completed using three datasets. Separate simulations were run for each herd and parasite species in each dataset. For all validation datasets, the per-Adult fecundity at the start of the simulations was set by estimating the level of acquired immunity (detailed below) and rearranging the equation for fecundity, *λ* (Table 1). This fecundity estimate was then divided by the initial faecal egg count (FEC) reported in the publications to estimate the initial number of Adults present, with an adjustment for daily faecal output included to ensure consistency.

First, the data used to validate the GLOWORM-PARA module in Rose Vineer et al. (2020; obtained from Verschave (2015), and referred to hereafter as the *Verschave* Herds 1-7), were used to validate the model for set-stocked cattle in Belgium in their first grazing season. The location of each herd in the Verschave dataset was estimated based upon nearest village or landmark reported in Verschave (2015) **(supplementary info)** and the *ncdf4* function v 1.22 in R (Pierce, 2019) was used to extract rainfall and temperature data from the E-OBS gridded dataset for each location (v 29.0e E-OBS (2023)). The standing biomass for each herd was available from Verschave (2015), and was estimated as 2000 kg DM for some herds where these data were not available. The initial level of acquired immunity, *r*, was set to values between 0.1 and 0.5 depending on the age of cattle in each herd at the start of simulations Vercruysse and Claerebout (1997). Initial pasture contamination levels were set to 0, with assumptions made that pastures were “clean” from parasite larvae, with just the initial number of Adult worms present within the host cattle.

Second, the model was validated for rotationally grazed cattle in their second grazing season using parasitological and farm management data obtained from four groups of cattle in Northern Ireland (McFarland et al., 2022; hereafter referred to as the *McFarland* Herds A-D). Faecal samples were collected twice monthly for FECs between turn out in May and housing in October. Cattle were rotated within their group-specific pastures (6 pastures for each group). It was possible to extract estimates of herbage biomass, movement dates, stocking rates, anthelmintic treatment dates and efficacy, approximate initial FEC, relative proportions of *O. ostertagi* and *C. oncophora*, and daily weather data from the publication. Based on descriptive detail in the publication, it was assumed that pasture was clean at the start of the simulation. Initial values for state variables were set to zero, except the number of Adults and the level of acquired immunity. The level of acquired immunity, *r*, was assumed to be 0.25 for *O. ostertagi* and 0.8 for *C. oncophora*, based on our current understanding of the development of protective immunity in calves over their first two grazing seasons (Ploeger et al., 1995), and an assumption that there is partial waning of immunity over the winter housing period.

Finally, we utilised data collected from experimentally infected animals grazed at Moredun Research Institute in Scotland (hereafter referred to as the “Passage herd”). Two parasite naive calves were infected with 20,000 *O. ostertagi* L3 (MOo2 isolate) and two calves with 20,000 *C. oncophora* L3 (MFie26 isolate). The pairs of calves infected with each parasite species were set-stock grazed on separate paddocks, neither of which had carried cattle prior to the study. Cattle were grazed May till October (18 weeks) and faecal egg counts were conducted at regular intervals to monitor parasite infection. Anthelmintic treatments in the empirical datasets were incorporated into simulations by removing the within-host population by a proportion consistent with the activity and efficacy of the anthelmintic product used.

Model performance against the validation datasets was evaluated by linear regression of the simulated and observed FECs, with an intercept forced through zero, as described previously (Rose Vineer et al., 2020; Rose et al., 2015; Wang et al., 2022). Because GLOWORM-META is a continuous mean-field model, it is considered useful if it is able to replicate the overall pattern of average FECs (assessed by visualisation of model output), and if the 95% confidence interval of the slope of the regression spans 1, where a slope of 1 indicates perfect model performance. As there is significant variation between individual FECs within herds which can be attributed to a multitude of factors (Coop and Kyriazakis, 2001; Milewski and Gümüş, 2023; Bricarello et al., 2023), including measurement error, *R*^2^ values are not expected to be high, but are nevertheless useful as they provide an indication of the proportion of variability in FECs that can be explained by a deterministic, climate-based model of the underlying population dynamics.

### 2.4. Evaluating GIN biosecurity strategies

#### 2.4.1. Sensitivity Analysis

The Quantities of Interest (QoIs) for model exploration of biosecurity factors are broadly divided into two factors affecting the safety of the resident herd: 1) the persistent contamination of quarantine areas up to 6 months after purchase of cattle (hereafter the “Residual contamination”), and 2) the additional exposure of the main herd to L3 contributed by purchased cattle after turnout (hereafter the “Additional exposure”).

To quantify the relative predicted importance of purchasing and quarantine parameters on these QoIs, the variance-based Sobol’ method for Global sensitivity analysis was applied to the model using the *soboljansen* function in the *sensitivity* package (Iooss and Weber, 2016), which generates samples (parameter combinations) for modelling and conducts Monte Carlo estimation of Sobol’ Indices on the model output. The Sobol’ Indices indicate the relative importance of the tested parameters by quantifying the model variance attributed to each parameter. First order indices indicate the variance in model output that can be attributed to each variable on its own. Total indices indicate the variance in model output that can be attributed to each variable, including higher order interactions with other variables.

The biosecurity parameters and model input of interest were: 1) duration of quarantine (1-90 days), 2) location of quarantine (housed/hard standing or on pasture), 3) date of purchase (day of year, indicating local seasonal/weather influences on the fate of eggs shed by surviving worms), 4) FECs at the time of purchase (0-900 eggs per gram), and 5) geographic location. The range of the duration of quarantine was based on a qualitative study of biosecurity practices on UK farms (N. Mahon & C. Hardy, unpublished data). The range of FECs at the time of purchase was based on cattle sampled in UK markets (L. Melville & D. Bartley, unpublished data). Geographic locations were randomly sampled using the *naturalearth* and *sf* packages (McGowan and O’Leary, 2021; Pebesma et al., 2023): 5,000 points within the United Kingdom (reflecting regional climatic influences) and 15,000 points across the wider European region (reflecting continental climatic influences). Each potential location was checked against the EOBS gridded dataset (v29.0e; (E-OBS, 2023)) and locations with no available data were rejected prior to sensitivity analyses (UK: n = 496; Europe: n = 1,717). Additionally, there were locations within Europe which had extreme weather which prevented simulation of the full life cycle. These locations were excluded to prevent later simulation errors (Europe: n = 5,250).

A GLOWORM-META simulation was conducted for each nematode species across sample scenarios for the UK (7,000) and Europe (24,000), representing parameter combinations generated by the *soboljansen* function. For each *O. ostertagi* simulation we assumed that cattle were 6 months of age at purchase, had initial resistance levels (*r*) of 0.25, and were stocked at 2 head per hectare. For *C. oncophora*, the same assumptions were made but the initial resistance was set at 0.5. The QoIs were derived from model output by extracting: 1) the total of L3 on the quarantine patch 6 months post-purchase, and 2) the sum of daily L3 on pasture between turnout of purchased calves with the resident herd and the end of simulations. Model output (the QoIs) was z-transformed before being returned *soboljansen* for estimation of the Sobol’ Indices and confidence intervals by bootstrapping (n = 100).

#### 2.4.2. Biosecurity Scenarios

Multiple simulations were developed to examine how the model would perform in practical biosecurity scenarios, simulating the movement of incoming cattle from markets to farms. This included scenarios with varying quarantine durations (7, 14, 30, 60 and 90 days), transitioning from the Quarantine pasture to a “Secondary” pasture. For the purposes of these scenarios, we began the simulations with no prior parasite burdens on either the quarantine or secondary pastures. Initial FECs within animals were set at three points, 50 eggs per gram (epg; representing low burdens typical of purchased cattle, L. Melville & D. Bartley, unpublished data), 500 epg (representing higher burdens observed in purchased cattle, L. Melville & D. Bartley, unpublished data), and at 900 epg (representing a very high burden). The hypobiosis of larval burdens was accounted for in the simulations through adjusting the initial partition of the parasite burden within the host to reflect seasonal differences. In simulated spring, a greater proportion of the initial parasite burden was allocated in the model to the Adult stage, while in simulated autumn, a higher proportion was allocated in the model to the Preadult stage to represent biological shifts in parasite development due to seasonal conditions. Anthelmintic treatments were simulated to occur on the day of arrival into the quarantine pasture (Day 1). Different treatment efficacy levels were used as a proxy for anthelmintic resistance: a 99% effective treatment represented good efficacy, an 80% effective treatment indicated moderate efficacy, and a 60% effective treatment reflected low efficacy. Acquired immunity levels were set to 0.01 for both *O. ostertagi* and *C. oncophora* to reflect young first-season grazer calves with no prior history of parasite burdens. The same geographic location as the Passage validation herds was used for these scenarios, with a herbage density of 2000 kgDM.

### 2.5. Larval mortality

To assess the duration of survival of infective L3 larvae on the quarantine pastures following cattle movement to secondary pastures, simulations were conducted to estimate the “half-life” of an initial L3 burden. The simulation was run for 100 randomly selected locations across the UK.

In each simulation, an initial burden of 100 L3 larvae was introduced to the pasture at the beginning of each month (i.e., on the first day of each month from January to December) using the *eventdat* argument in the *lsoda* function v 1.24 (Soetaert et al., 2010). This process was therefore repeated each month for the year 2020 to assess the influence of seasonality on L3 survival. Simulations were performed using the GLOWORM-META model, under the assumption that there were no cattle were present (i.e., a stocking rate of zero) and with a standardised pasture biomass of 2000 kg DM. Each simulation was conducted separately for both *Ostertagia* and *Cooperia* parameter sets.

Following the simulations, the “half-life” of L3 was calculated for each location and month, defined as the number of days required for the initial burden of 100 L3 larvae to decline to 50 L3. To assess temporal variation in the half-life duration over months, Generalized Additive Models (GAMs) (using the *gam* function in the *mgcv* package (Wood, 2017)), were applied separately for *Ostertagia*and *Cooperia* species. These models allow for an understanding of how the half-life varied by month and determine whether there were significant temporal patterns in L3 survival. The GAM models were also used to assess the influence of environmental factors, such as the temperature and precipitation, on the half-life duration for each species. Violin plots were generated (using the *vioplot* function within the R package *vioplot* (Adler et al., 2023)), to visualize the distribution of L3 survival times across different locations and months. To assess temporal variation in the half-life duration of *O. ostertagi* and *C. oncophora* across months, Kruskal-Wallis rank sum tests were performed separately for each species. Subsequent pairwise comparisons were conducted using the Wilcoxon rank sum test with continuity correction to identify which specific months differed significantly. These pairwise tests were adjusted for multiple comparisons using the Bonferroni correction. For clarity, only p-values less than 0.001 (*p <* 0.001) were considered statistically significant and are reported in the results.

### 2.6. Ethical approvals

The data from the “Passage herd” and the data used to estimate vertical migration rates, both described above, were collected using excess faecal material generated during the routine passage of isolates at the Moredun Research Institute. This was approved by the Moredun Research Institute’s Animal Welfare Ethical Review Body (AWERB), 20th January 2023; project title: Control of nematodes and anthelmintic resistance, experiment number: E05/23, Home Office Project Licence No. PP6939295, Personal Licence No. 14802EC9C. Secondary use of the data for model development was approved by University of Liverpool’s AWERB 29th May 2024, approval No. AWC0280.

## 3. Results

### 3.1. Validation

The model, parameterised for both species, produced plausible patterns and magnitudes of FECs over the course of a grazing season for the majority of herds, with individual FECs deviating from the mean simulations as expected (Figure 2 and 3). Linear modelling confirmed good overall concordance between observed and simulated FECs (*O. ostertagi*; slope = 1.36 (0.117 SE), *C. oncophora*; slope = 1.53 (0.101 SE), although unsurprisingly, unexplained variability in the FECs remained (*O. ostertagi*; *R*^2^ = 0.234, *C. oncophora R*^2^ = 0.344), where the model captured mean FECs but not the high individual FECs in the observed data (Figure 2 and 3, and the Supplementary figures A.11 and A.12).

**Figure 2:**
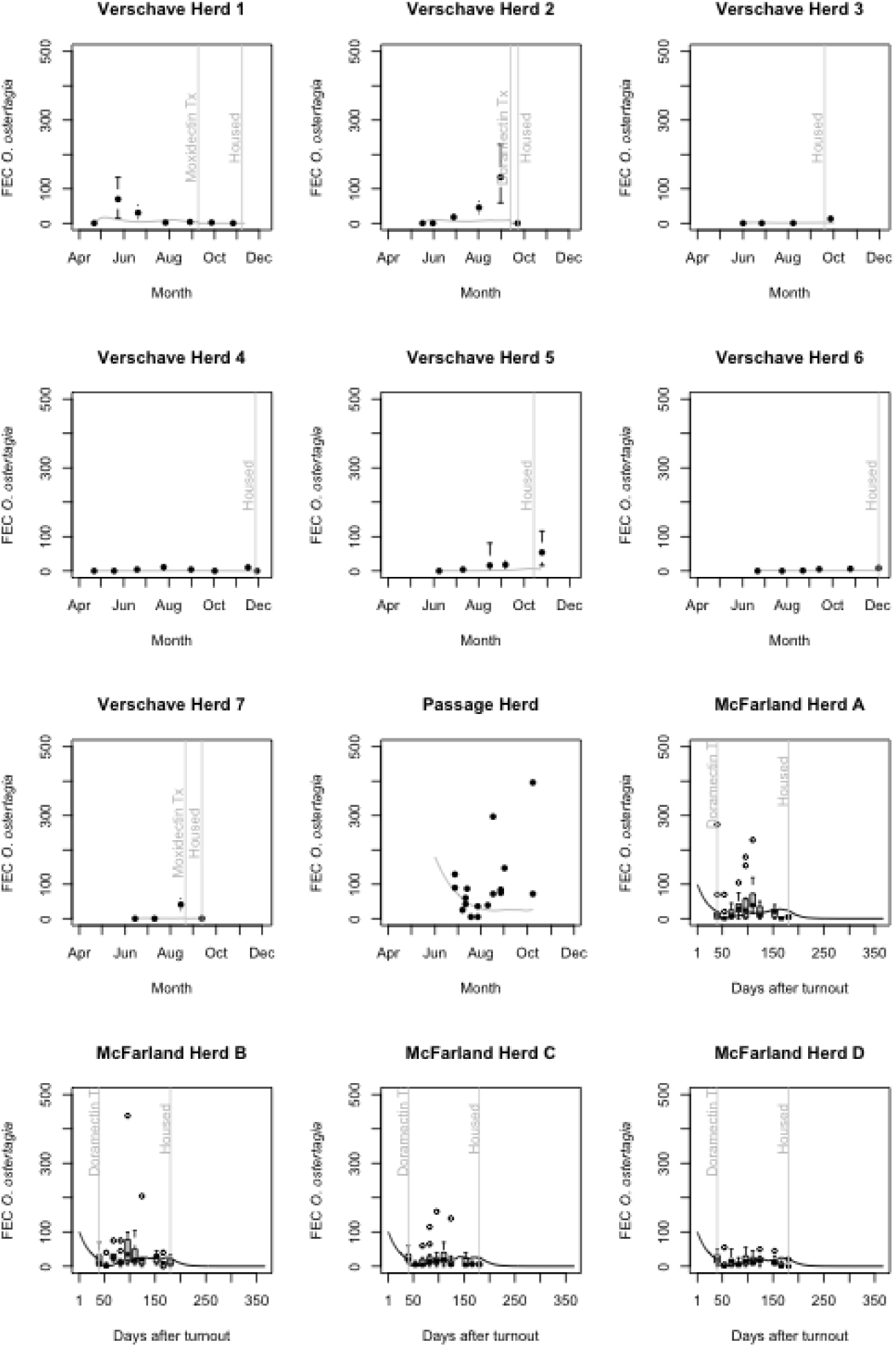
Predicted and observed faecal egg counts for *O. ostertagi* for the GLOWORM-META validations; single pastures (Verschave et al, 2015; Moredun Passage Herd), and with multiple pasture rotations (McFarland et al, 2021). Further information on the background of the data used can be found in the supplementary information. The points and error bars show the number of eggs per gram, and the 95 confidence interval around the mean. The solid grey line depicts the simulated faecal egg counts. Light grey vertical ablines represent key management events in the empirical datasets which were also included in model simulations.

**Figure 3:**
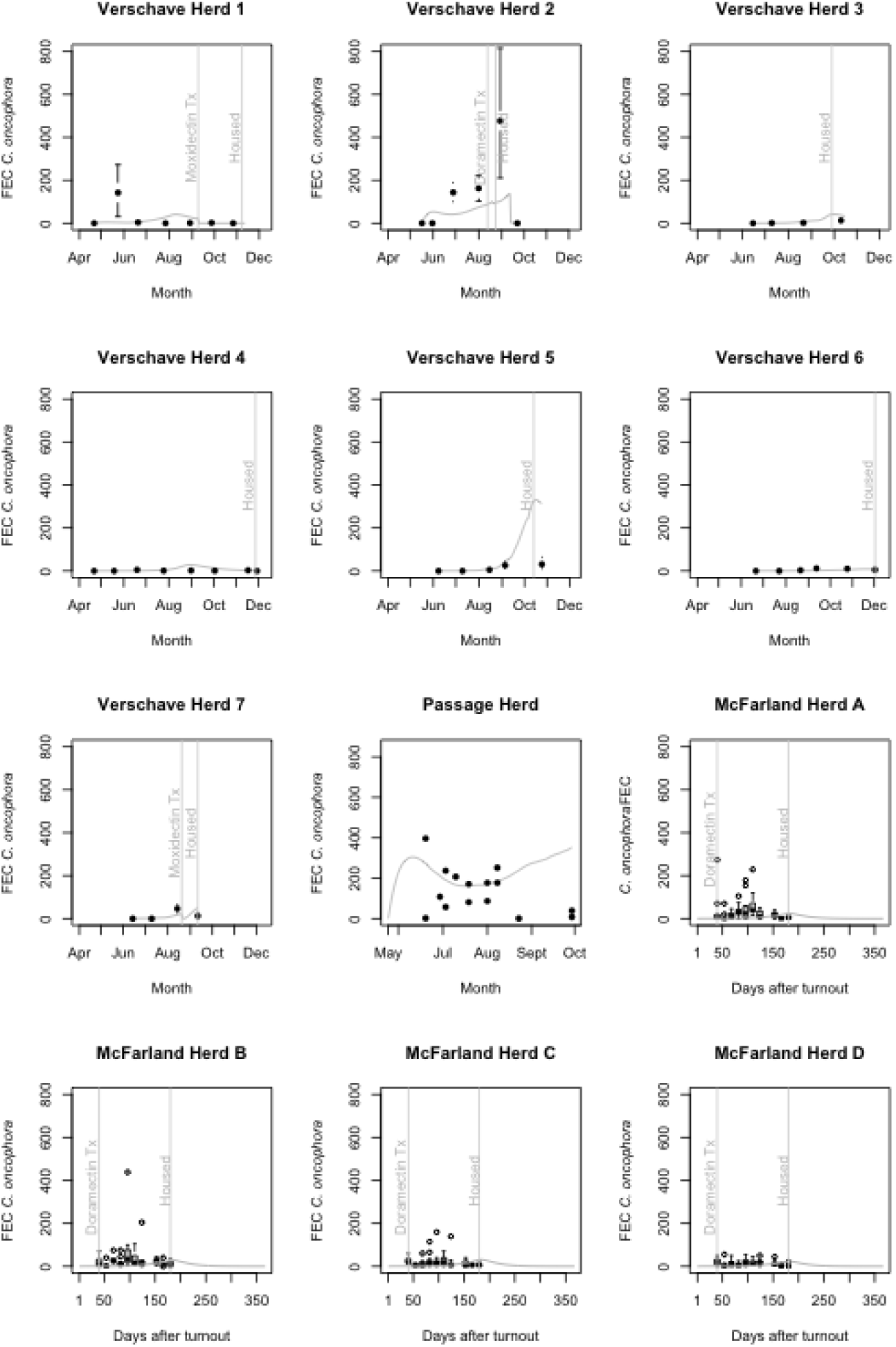
Predicted and observed faecal egg counts for *C. oncophora* for the GLOWORM-META validations; single pastures (Verschave et al, 2015; Moredun Passage Herd), and with multiple pasture rotations (McFarland et al, 2021). Further information on the background of the data used can be found in the supplementary information. The points and error bars show the number of eggs per gram, and the 95 confidence interval around the mean. The solid grey line depicts the simulated faecal egg counts. Light grey vertical ablines represent key management events in the empirical datasets which were also included in model simulations.

### 3.2. Sensitivity analyses

Global sensitivity analysis results were largely consistent across both the UK and European geographic datasets, and across species (Figure 4 and 5). The duration of quarantine emerged as the most influential parameter for additional exposure to the resident herd (in the absence of quarantine anthelmintic treatment), with high first order (main effect) and total (total effect) SIs. Total SIs were higher than the first order SIs, indicating interacting effects with other parameters.

**Figure 4:**
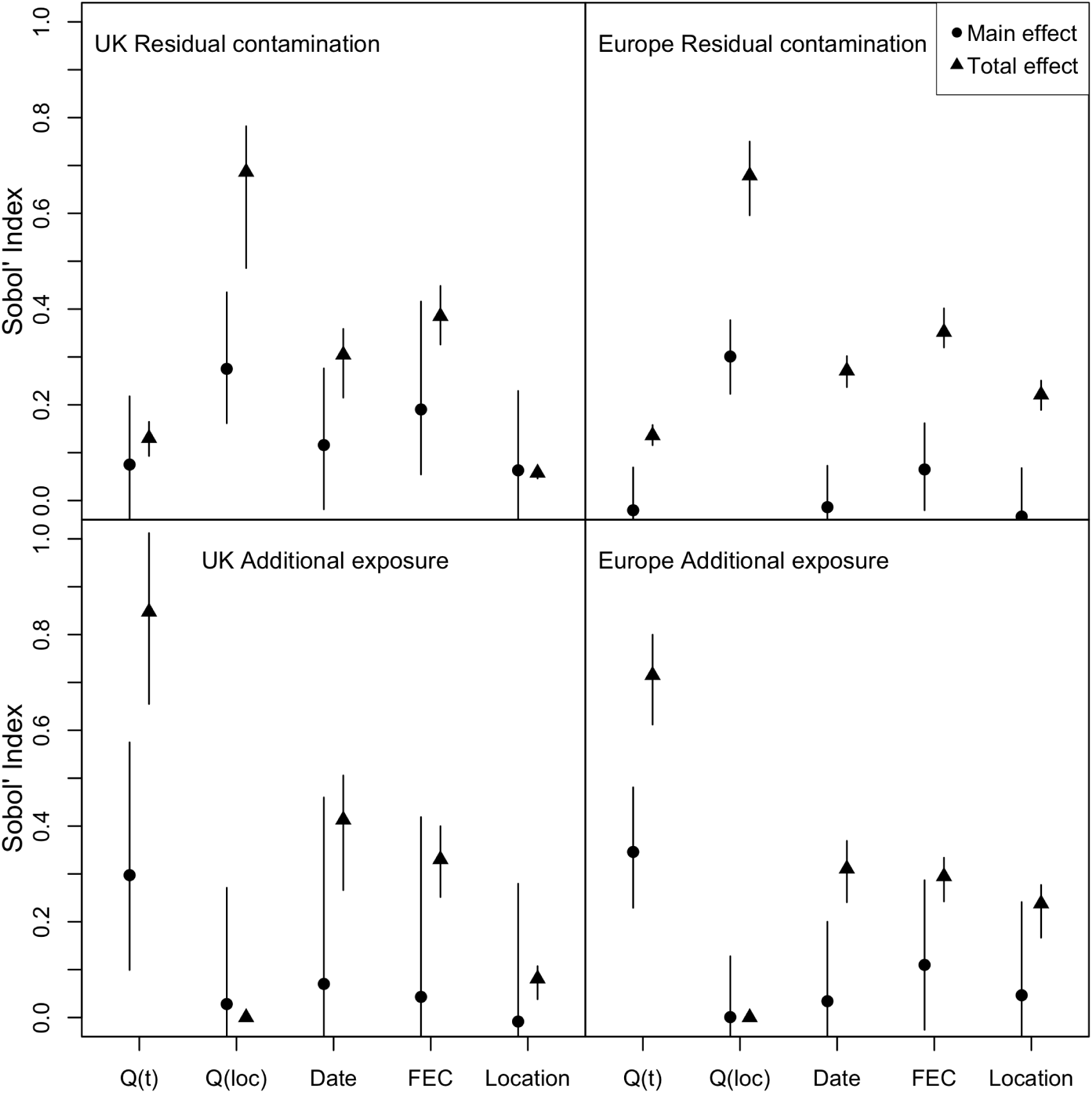
Sobol’ Indices and 95% bootstrapped confidence intervals for *O. ostertagi* sensitivity analyses using geographic locations from the UK and Europe. The five model parameters tested (x-axis; Q(t) = quarantine time (days), Q(loc) = quarantine location (pasture or housed/yarded), Date = purchase date (day of year), FEC = faecal egg counts at the time of purchase (eggs per gram), Location = geographic location of the simulated purchasing farm) were varied and evaluated against two Quantities of Interest (Residual contamination = total L3 on pasture remaining at the end of the simulations. Additional exposure = sum of the daily L3 on pasture contributed by purchased cattle after joining the main herd (time dependent on the length of quarantine allocated in the residual contamination, between 0-90 days following, and until 6 months after the purchase date), representing the additional exposure of the resident herd to GINs contributed by the purchased cattle). Round points indicate the main effect (first order Sobol’ Index) and triangles represent the total effect including all higher order interactions.

**Figure 5:**
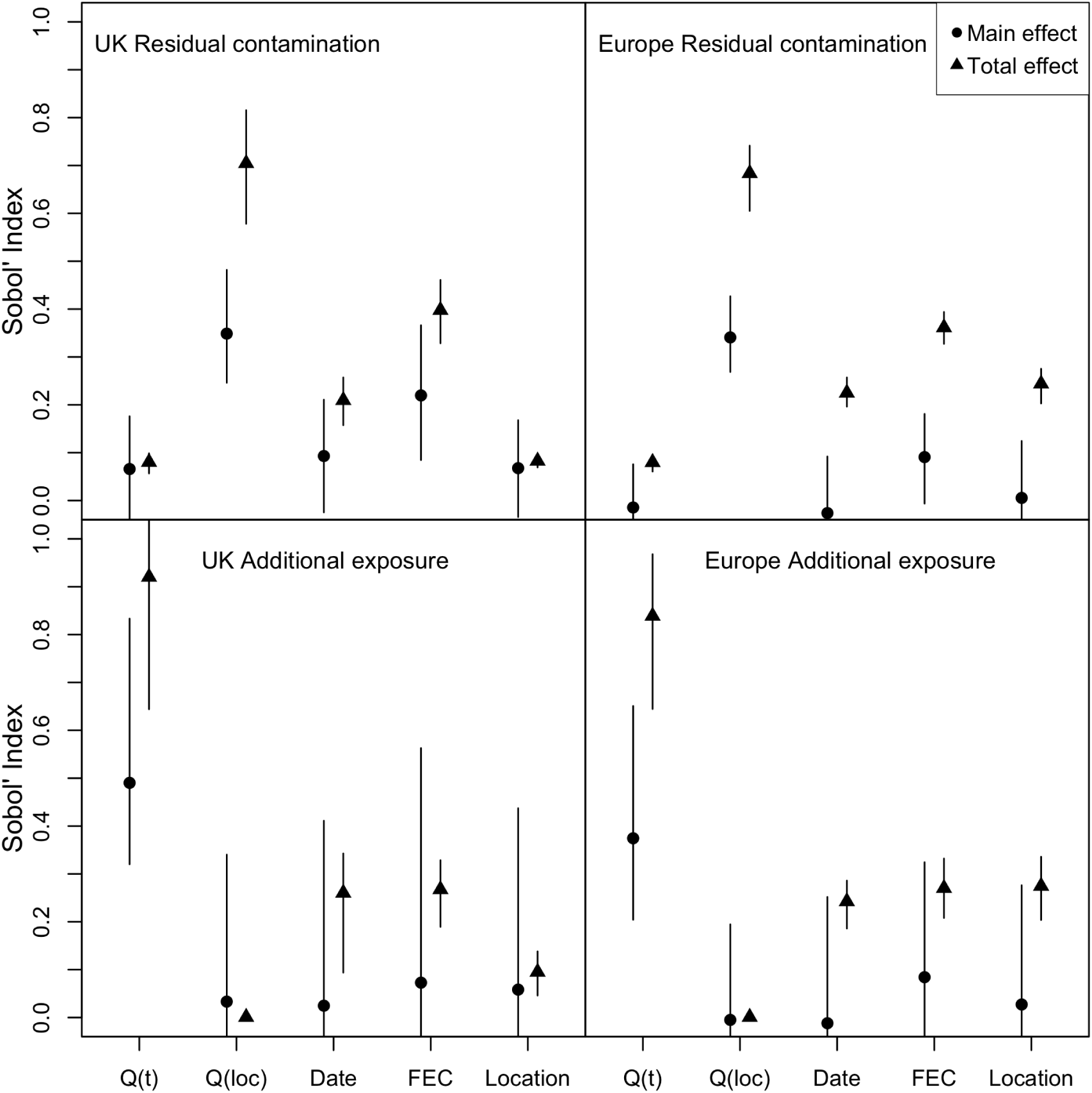
Sobol’ Indices and 95% bootstrapped confidence intervals for *C. oncophora* sensitivity analyses using geographic locations from the UK and Europe. The five model parameters tested (x-axis; Q(t) = quarantine time (days), Q(loc) = quarantine location (pasture or housed/yarded), Date = purchase date (day of year), FEC = faecal egg counts at the time of purchase (eggs per gram), Location = geographic location of the simulated purchasing farm) were varied and evaluated against two Quantities of Interest (Residual contamination = total L3 on pasture remaining at the end of the simulations. Additional exposure = sum of the daily L3 on pasture contributed by purchased cattle after joining the main herd (time dependent on the length of quarantine allocated in the residual contamination, between 0-90 days following, and until 6 months after the purchase date), representing the additional exposure of the resident herd to GINs contributed by the purchased cattle). Round points indicate the main effect (first order Sobol’ Index) and triangles represent the total effect including all higher order interactions.

In both the UK and Europe, the date of purchase (influencing weather patterns during the simulations) and FECs on the day of purchase were also moderately influential on both QoIs when considering interactions with other parameters (total effect). Geographic location had minimal influence on model variance for the residual contamination and additional exposure QoIs in the UK but did influence the two QoIs in Europe.

A large proportion of variability in the residual contamination of quarantine pastures was attributed to the location of quarantine (Q(loc)) (Figure 4 and 5), which was expected as housing or yarding would generate zero residual contamination when compared to housing on pastures. The location of quarantine (on hard standing/housed or on pasture) thus yielded high QoIs for the residual contamination. Quarantine location was less influential on additional exposure.

Visualisation of the residual contamination and additional exposure QoIs against the duration of quarantine, farm latitude (UK), purchase date and FECs at purchase provides additional information about the pattern of influence of the parameters tested (Figure 6 and 7). Additional exposure decreased rapidly with increasing quarantine duration, and residual contamination increased with duration of quarantine, though with greater variability. Variability of both QoIs was influenced by date of purchase, increasing in the first half of the year and decreasing thereafter. Variability in both QoIs increased with increasing FEC on the day of purchase. Consistent with the Sobol’ Indices, there was no apparent pattern to the distribution of QoIs plotted against purchasing farm latitude within the UK. However, in the European Sobol Indices, a subtle pattern emerged in the QoI distribution for the geographic location. This was similarly reflected in the QoIs plotted against purchasing farm latitude (Figure 8 and 9).

**Figure 6:**
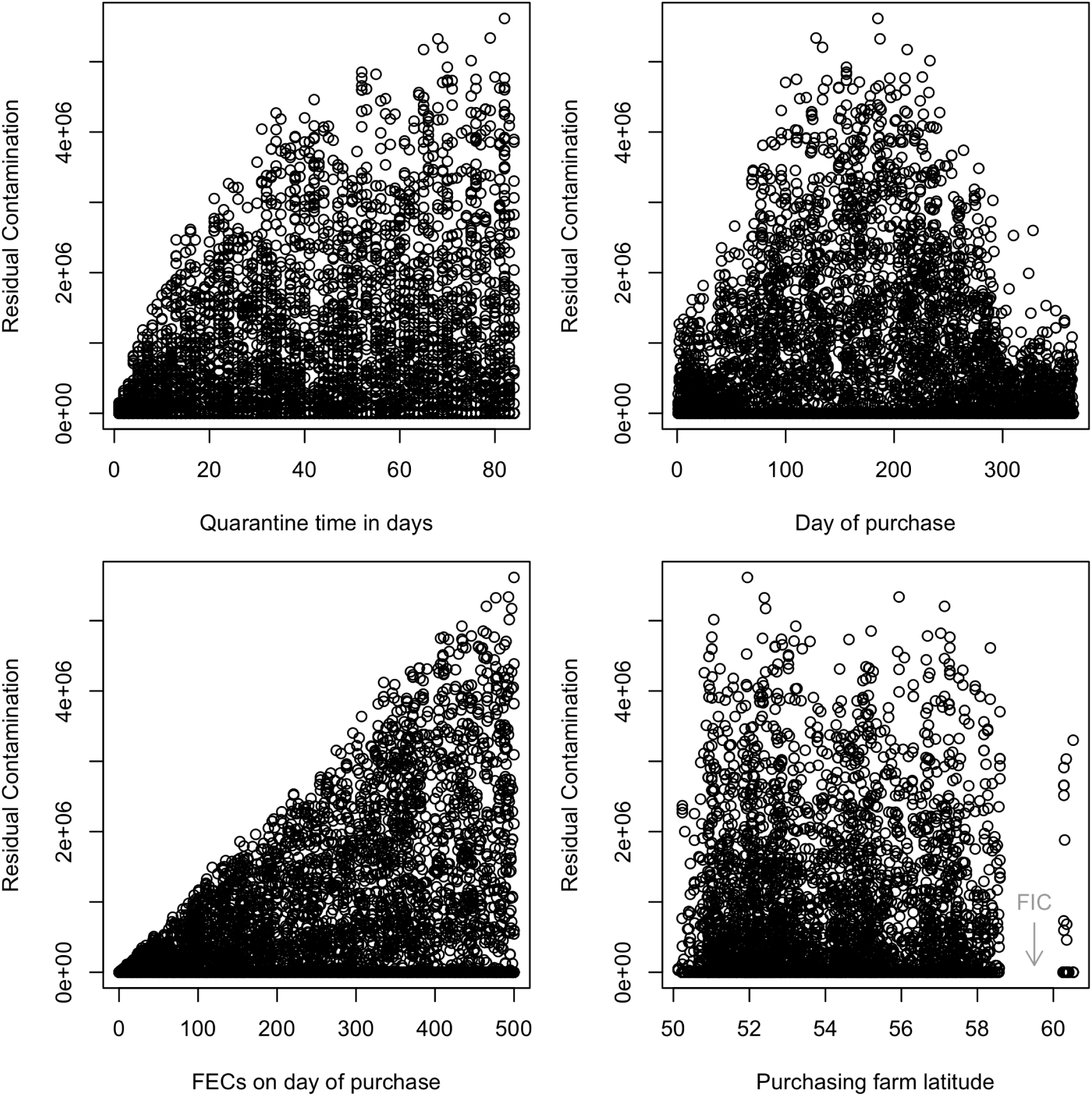
Residual Contamination for *O. ostertagi* simulations in the UK against the four parameters: 1) Quarantine time in days, 2) Day of purchase (where day 1 corresponds to the 1st January), 3) FECs on the day of purchase, 4) the purchasing farm latitude. FIC = Fair Isle Channel.

**Figure 7:**
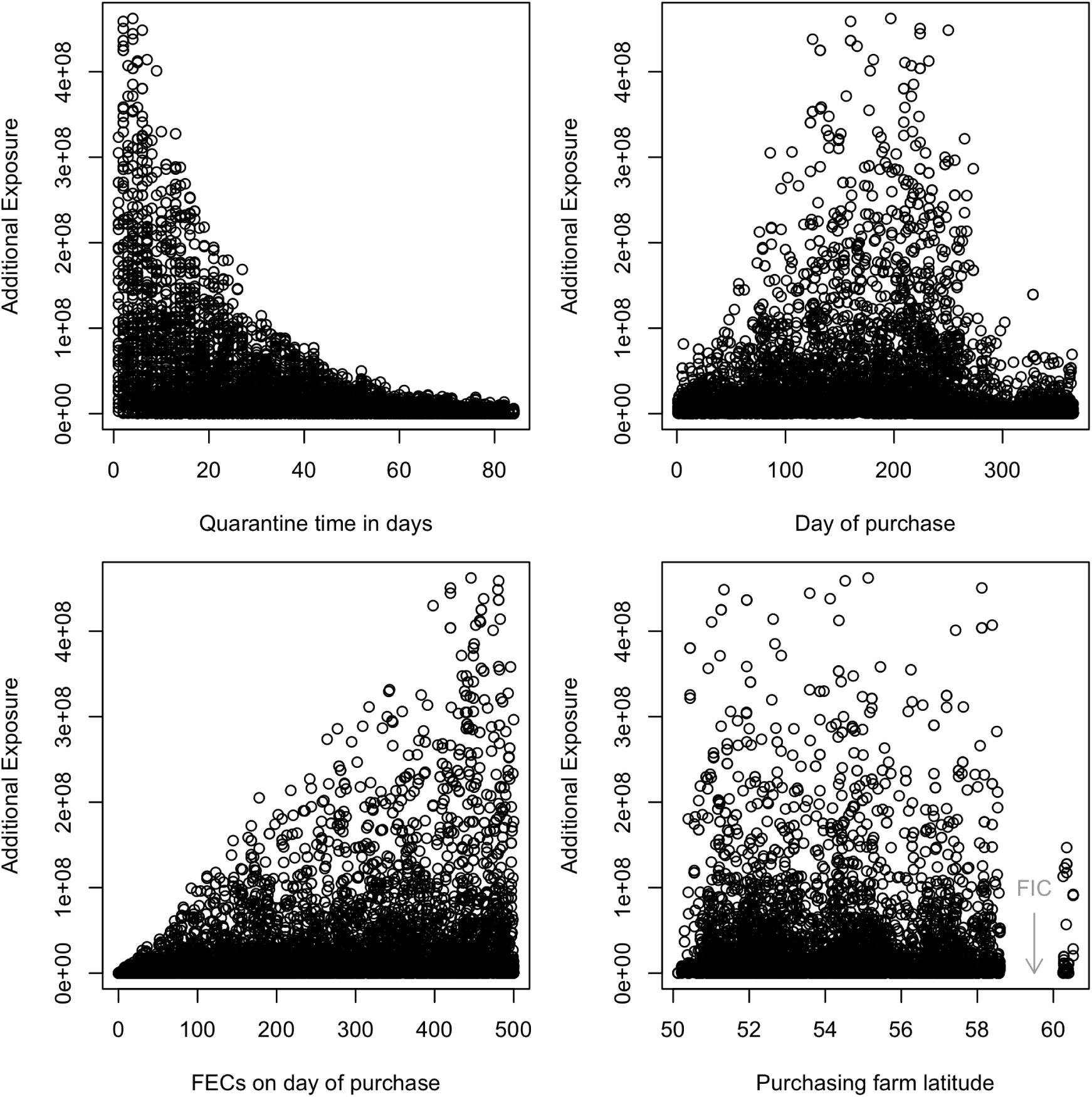
Additional Exposure for *O. ostertagi* simulations in the UK against the four parameters: 1) Quarantine time in days, 2) Day of purchase (where day 1 corresponds to the 1st January), 3) FECs on the day of purchase, 4) the purchasing farm latitude. FIC = Fair Isle Channel.

**Figure 8:**
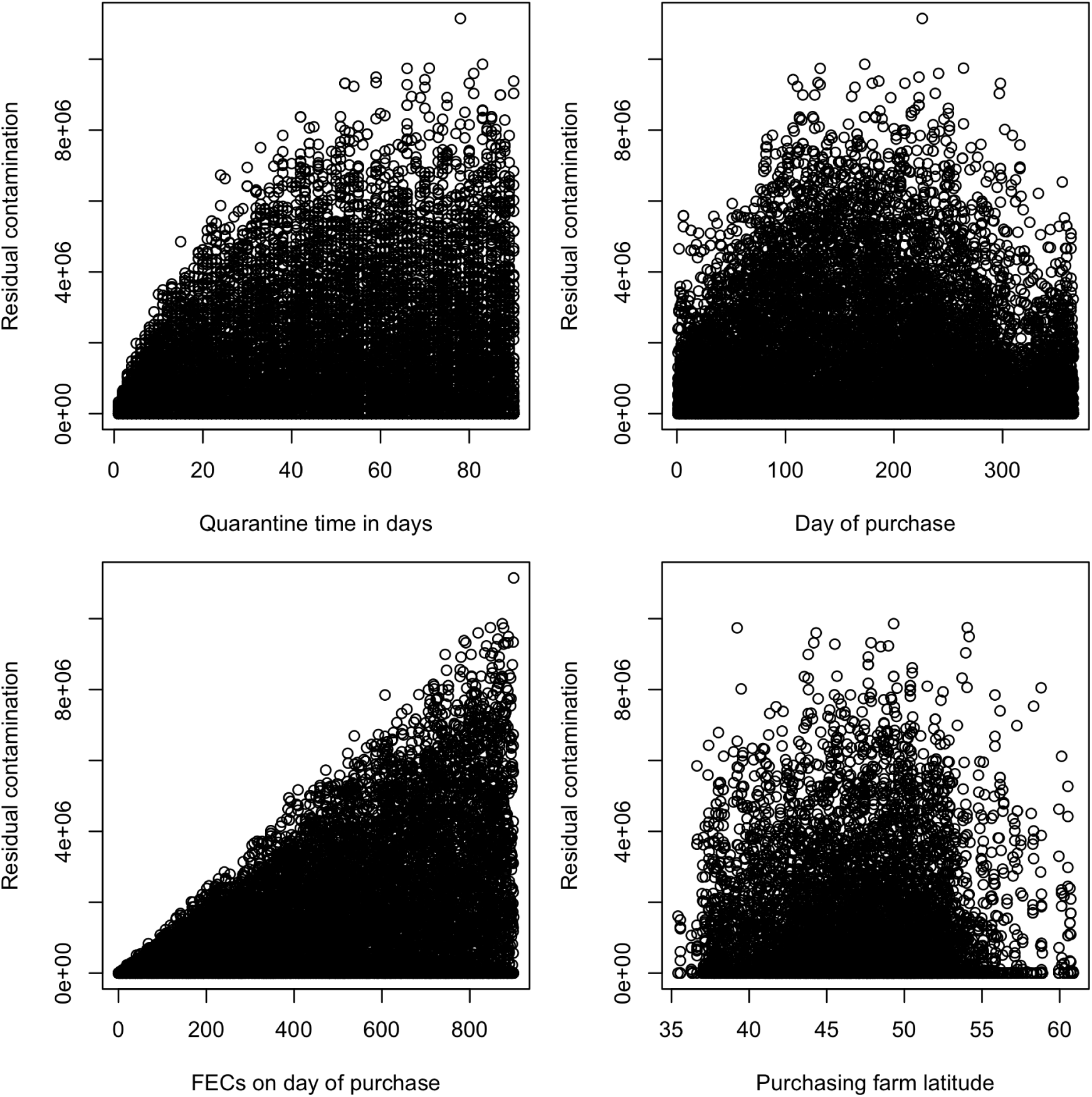
Residual Contamination for *O. ostertagi* simulations in Europe against the four parameters: 1) Quarantine time in days, 2) Day of purchase (where day 1 corresponds to the 1st January), 3) FECs on the day of purchase, 4) the purchasing farm latitude.

**Figure 9:**
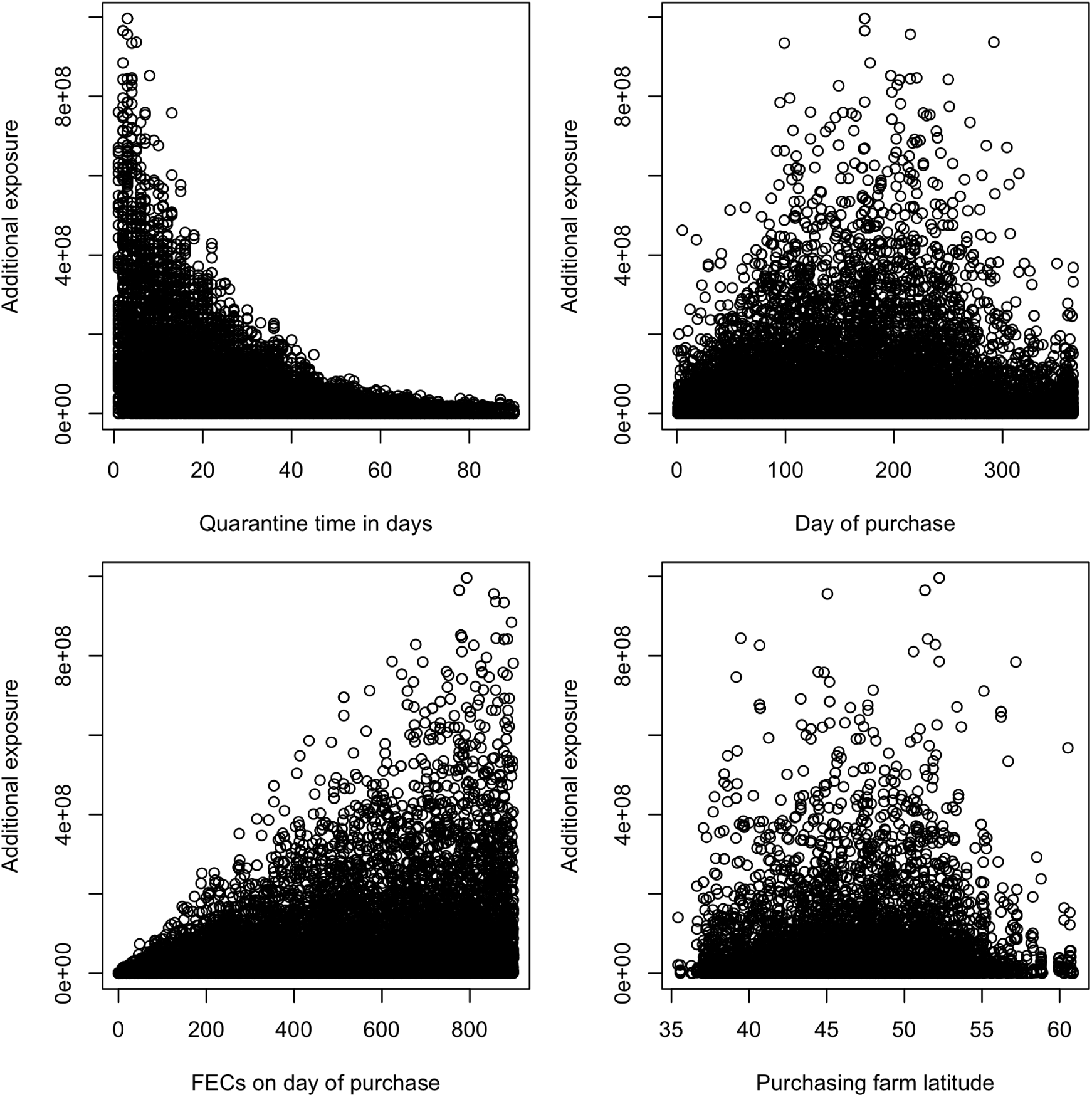
Additional Exposure for *O. ostertagi* simulations in Europe against the four parameters: 1) Quarantine time in days, 2) Day of purchase (where day 1 corresponds to the 1st January), 3) FECs on the day of purchase, 4) the purchasing farm latitude.

### 3.3. Biosecurity Scenarios

The simulations revealed realistic patterns of infective larvae (L3) distribution across the modelled pastures: the Quarantine (initial) pasture (L3p A) and the Secondary pasture (L3p B)(Figure 10). In all scenarios, animals initially grazed on the quarantine pasture before transitioning to the secondary pasture.

**Figure 10:**
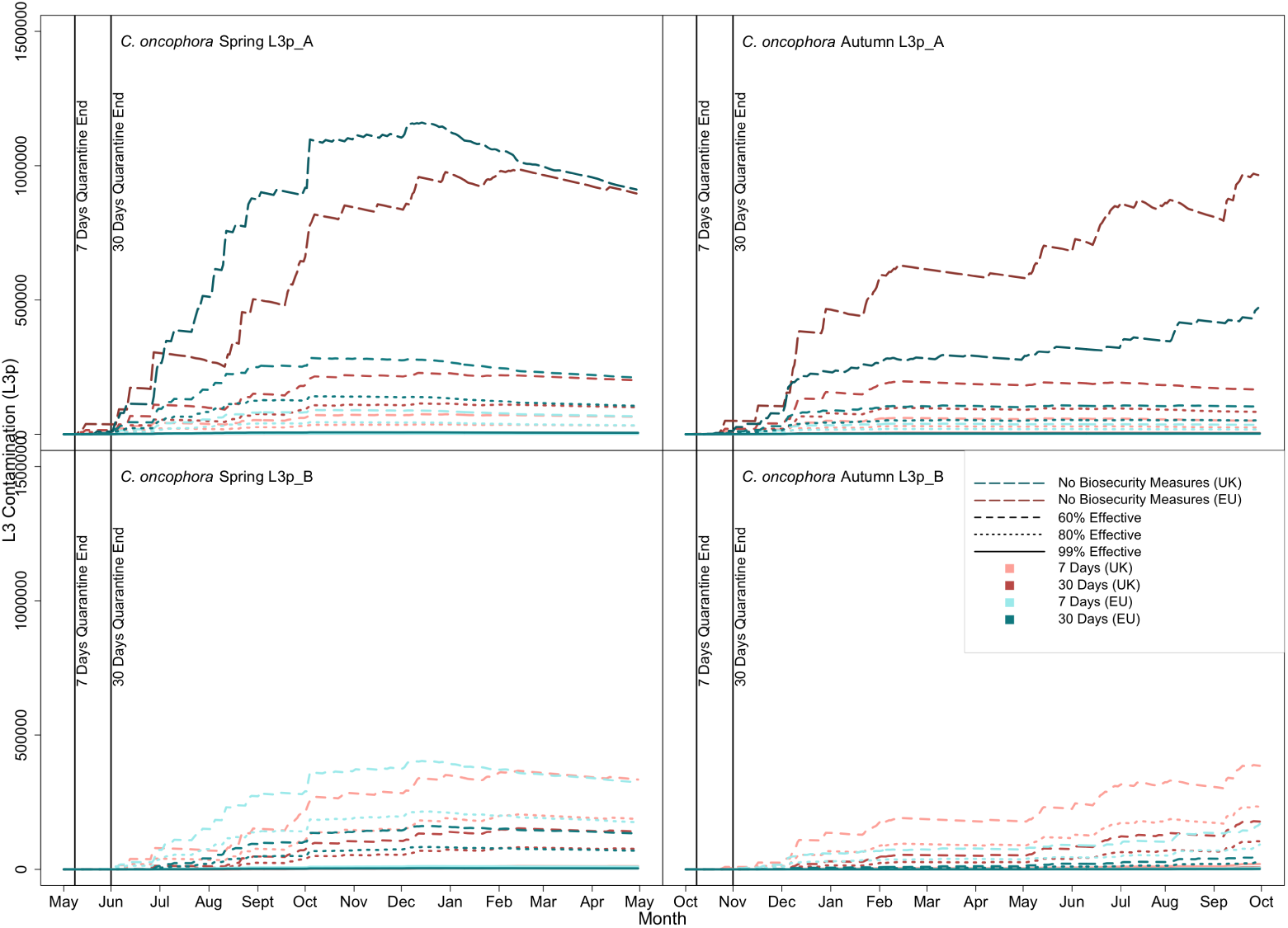
Biosecurity simulation for an initial FEC of 500 epg for *C. oncophora*. Comparison of the UK (Penicuik, Scotland) and southern France (Cahors, France) for both spring (May) and autumn (October) purchase. These graphs show the L3 contamination for the Quarantine pasture (L3p A) and the secondary pasture (L3p B) following differing lengths of quarantine and treatment interventions. Darkest lines represent no biosecurity measures (no treatments and no pasture movements). The effectiveness of anthelmintics represents resistance: 99% effective = little to no resistance, and 60% effective = high resistance.

For both parasite species, longer quarantine periods led to increased L3 contamination on the quarantine pasture and decreased L3 contamination on the secondary pasture. Furthermore, treatments with higher efficacy (used as a proxy for reduced anthelmintic resistance) resulted in lower L3 contamination levels on both pastures. To account for hypobiosis, the animals’ initial parasite burden (10,000 larvae per gram of feces) was adjusted for seasonal differences (Figure 10 and 11). During summer, this burden was distributed as 8,000 larvae in the adult stage and 2,000 in the preadult stage. In winter, the distribution shifted to 6,000 adult-stage larvae and 4,000 preadult-stage larvae.

**Figure 11:**
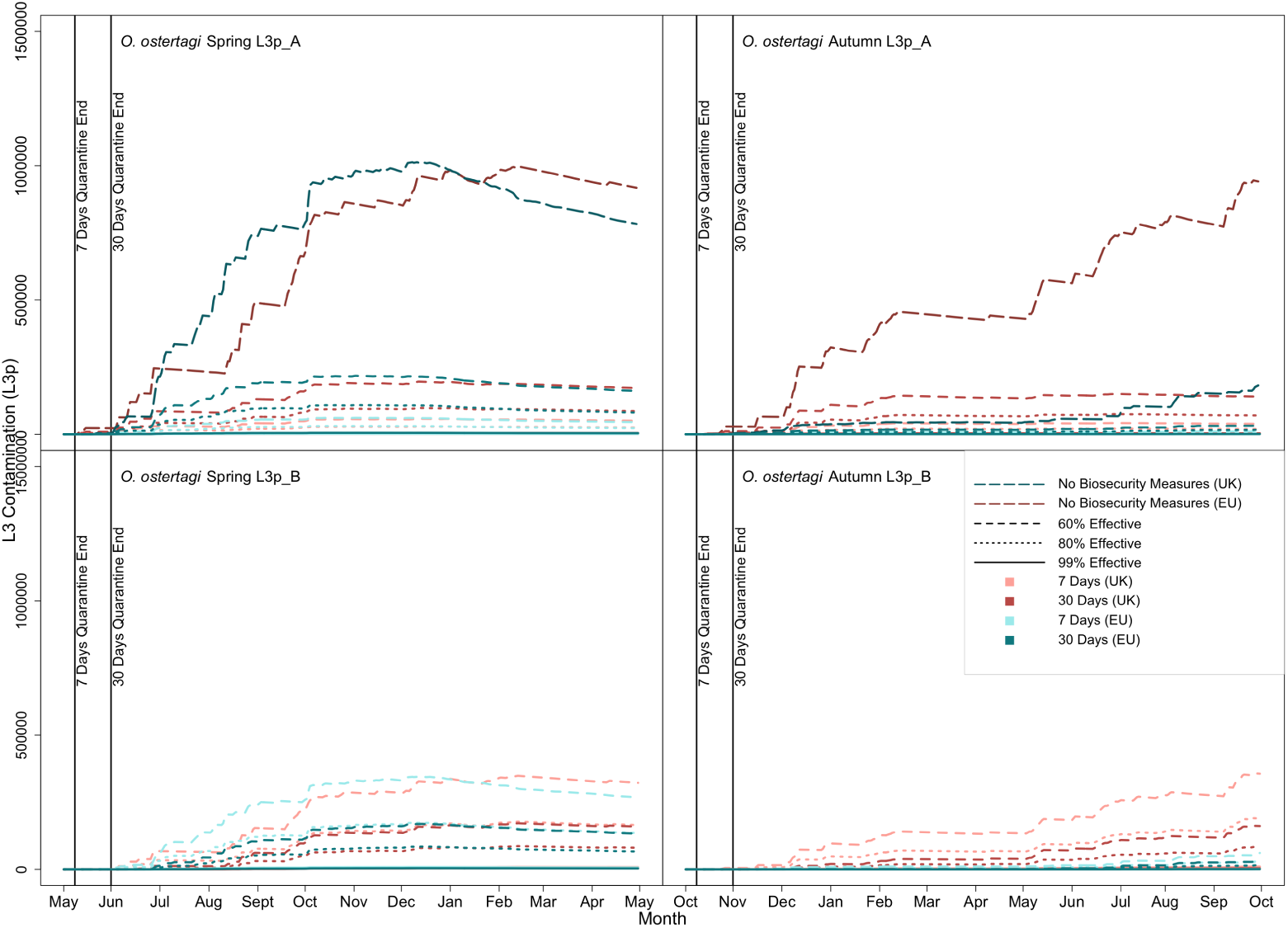
Biosecurity simulation for an initial FEC of 500 epg for *O. ostertagi* for comparison of the UK (Penicuik, Scotland) and southern France (Cahors, France) for both spring (May) and autumn (October) purchase. These graphs show the L3 contamination for the Quarantine pasture (L3p A) and the secondary pasture (L3p B) following differing lengths of quarantine and treatment interventions. Darkest lines represent what would occur if no biosecurity measures were used (no treatments and no pasture movements), where cattle were put straight out to pasture. The effectiveness of anthelmintics represents anthelmintic resistance, where 99% effective indicates little to no resistance, and 60% effective indicates high resistance.

### 3.4. Larval mortality

The time for the initial L3 burden (100 L3p) to drop below 50% (50 L3p) varied across months in the UK. Generalized Additive Models (GAMs) were applied separately for *O. ostertagi* and *C. oncophora* to examine significant differences in larval mortality. Both species exhibited significant relationships between larval half-life and month of the year (*O. ostertagi*: F = 119.462, *p <* 2*e −* 16, R² adj = 0.916; *C. oncophora*: F = 32.980, *p <* 2*e −* 16, R² adj = 0.858). However, the seasonal effect was more pronounced in *O. ostertagi* (Figure 12). Whilst it is recognised that they were used to simulate the model, temperature and precipitation were significant predictors of half-life duration for both species (*O. ostertagi*: Temperature: F = 1071.565, *p <* 2*e −* 16, R² adj = 0.916; Precipitation: F = 74.791, *p <* 2*e −* 16, R² adj = 0.916; *C. oncophora*: Temperature: F = 671.042, *p <* 2*e −* 16, R² adj = 0.858; Precipitation: F = 86.762, *p <* 2*e −* 16, R² adj = 0.858).

**Figure 12:**
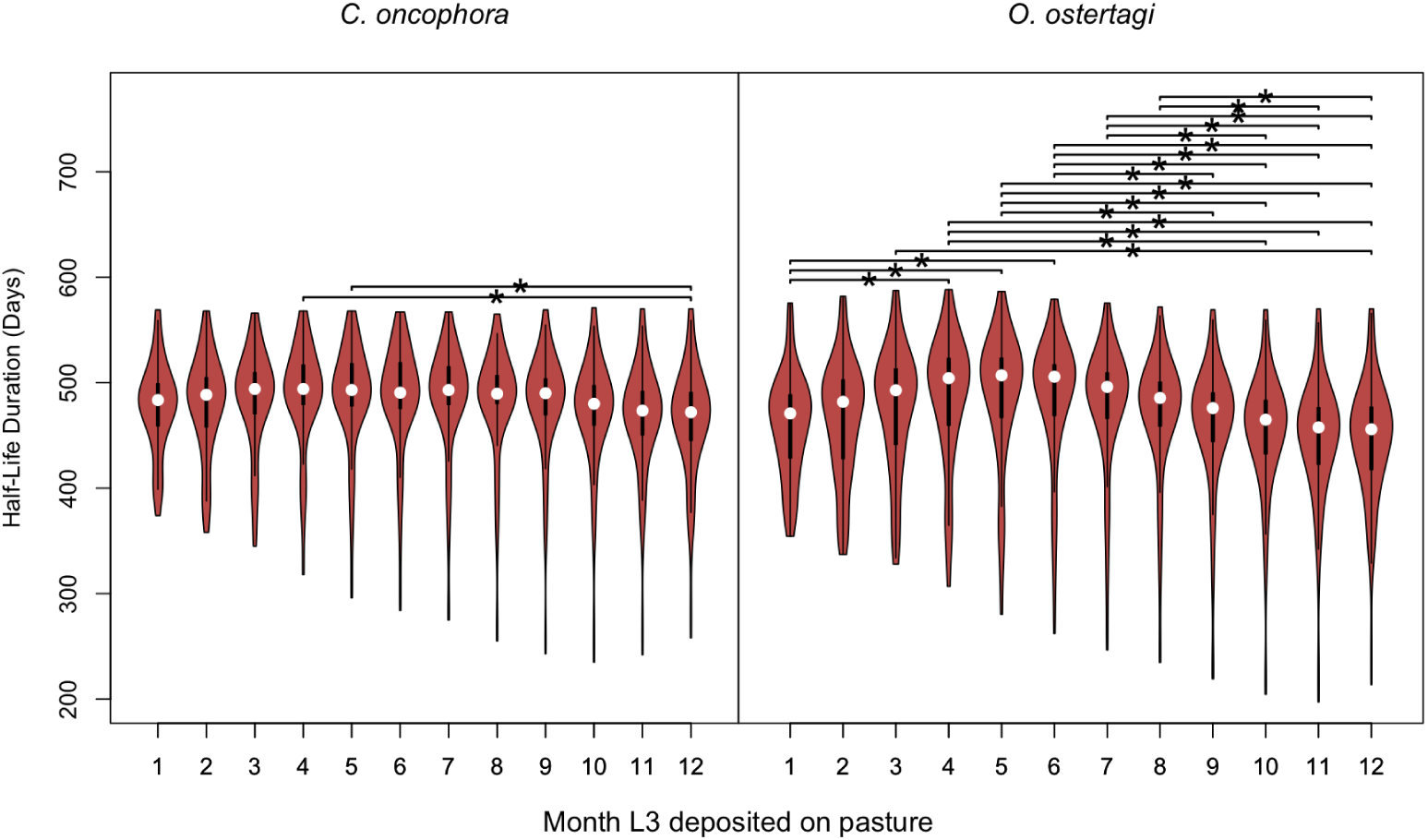
The half-life of an initial burden of 100 L3 larvae on pasture across 100 locations in the UK. Simulations were run monthly to assess seasonal variations in larval mortality. Month 1 represents January, through to Month 12 (December). The seasonal mean half-life durations with standard errors were as follows: for *O. ostertagi*, winter = 481.54 days (± 3.48), spring = 510.15 days (± 3.87), summer = 504.69 days (± 3.85), and autumn = 478.21 days (± 3.70); for *C. oncophora*, winter = 474.01 days (± 2.79), spring = 486.79 days (± 2.87), summer = 485.24 days (± 3.02), and autumn = 473.25 days (± 3.08). Significant differences (*p <* 0.001) between months are indicated by *.

For *O. ostertagi*, the average larval half-life increased from January to June, reaching its peak in late spring and early summer, before declining in the later months. Significant differences (*p <* 0.001) were found between several months. In contrast, *C. oncophora* showed a relatively stable half-life duration throughout the year, with fewer seasonal fluctuations. Seasonal mean half-life durations differed between species, with *O. ostertagi* having the longest mean duration in spring (510.15 days), and *C. oncophora* also peaking in spring (486.79 days) (Figure 12).

## 4. Discussion

A novel full life-cycle model of cattle GIN metapopulation dynamics was developed to proactively explore potential biosecurity measures which minimise introduction of GINs onto cattle enterprises. Previous cattle nematode transmission models tended to focus on only *O. ostertagi*. This is likely due to the pathogenicity of *O. ostertagi* compared to *C. oncophora*, practical difficulties differentiating mixed-species infections, and fewer sources of published data for *C. oncophora* (Verschave et al., 2016). However, advances in field data analyses have allowed parameterisation and validation of the GLOWORM-FL model for *C. oncophora* (Wang et al., 2022), which was further extended to the parasitic stages module in this study. The GLOWORM-META model, therefore, offers a unified framework for deterministic simulation of GIN metapopulation dynamics for both *Ostertagia ostertagi* and *Cooperia oncophora*, incorporating realistic farm management practices such as rotational grazing, treatments, and housing.

Global sensitivity analyses suggest that the influence of seasonal weather patterns on the relative impact of some GIN management strategies may outweigh that of regional climate differences at a national scale, but climate differences become similarly important at a continental scale. The influence of weather and climate on patterns of GIN transmission has received increasing attention within the veterinary parasitology research community due to the considerable impact of these factors on GIN phenology (van Dijk et al., 2008) and the potential to harness climate- and weather-driven variability to better target GIN management approaches (Gethings et al., 2015). For example, understanding the seasonal patterns of cattle GIN transmission may assist purchasers in making dynamic risk assessments that inform their grazing management, in cases where quarantine treatment has been ineffective.

Geographic location had a negligible influence on simulation variation in the UK in this study, but demonstrated an influence on the European simulations. This is likely attributed to the wider range of biogeographical regions, and associated heterogeneous climate, throughout Europe, spanning Northern boreal and Alpine regions (e.g., Norway) to southern Mediterranean regions (e.g., Spain). In contrast, in both the UK-only and the European-wide analyses, the date of purchase of calves (seasonality effect) ranked highly in the sensitivity analyses. The additional exposure of the resident herd to L3 peaked when calves were purchased mid-year and was lowest during winter, reflecting general seasonal temperature patterns and the theoretical impact of temperature on the success of GIN development Rose et al. (2015); Bolajoko et al. (2015).

The negligible influence of geographic location on simulation outcomes within the UK suggests that regional variations in biosecurity recommendations within a nation are not warranted without further supporting evidence. However, at broader spatial scales across Europe, significant climatic differences between regions may influence the outcomes of gastrointestinal nematode (GIN) control measures (Barnes and Dobson, 1990; Dobson et al., 2011). There is also variability in agricultural practices between countries. As a result, biosecurity recommendations may benefit from tailoring to the specific climatic, seasonal patterns and agricultural practices of each country or region.

The moderate-high influence of FECs at the time of purchase on the outcomes of interest in this study highlights the benefit of treating purchased cattle against GIN infections. However, with resistance to some anthelmintic classes observed on an average of 30% of European cattle farms, and an overall trend for increasing levels of anthelmintic resistance over time Taylor (2012); Kelleher et al. (2020); Rose Vineer et al. (2020a), purchasers should be cognisant of the possibility of treatment failure, and the need to minimise selection for anthelmintic resistance. Thus, although treatment failure was not explicitly simulated in the global sensitivity analysis, the range of FECs included in simulations implicitly represents potential populations of resistant GINs introduced onto a farm in the event of anthelmintic resistance, as demonstrated in the example simulations. These demonstrate how ineffective treatments would theoretically result in greater contamination of pasture contributed by the purchased cattle and their “resistant” nematodes. The benefits of treating purchased cattle are therefore balanced against the risks of selecting for anthelmintic resistance. Industry recommendations for GIN control in purchased cattle include anthelmintic treatment and post-treatment efficacy checks. Additional measures mitigating against the introduction of resistant genotypes should also be considered (Hoste and Torres-Acosta, 2011), such as increasing the duration of quarantine and re-treatment with a second anthelmintic class where post-treatment checks indicate treatment failure.

Quarantine pastures are often the heavily-used easily-accessible pasture, and careful recommendations on quarantine duration and management of higher burdens in quarantined areas are therefore essential. In our analyses, quarantine time, for both QoIs, and quarantine location, for the residual contamination QoI, were the most influential parameters of those tested. Longer simulated quarantine times reduced parasite burdens in secondary pastures (additional exposure QoI) but increased contamination within quarantine pastures (residual contamination QoI). Previous research has shown a reluctance among farmers to allocate or reserve specific space to hold quarantined animals, due to the cost and availability of appropriate areas to do this within farms (Damiaans et al., 2018). Therefore, although extending quarantine as much as possible allows additional time for existing infections (or anthelmintic resistant GINs remaining after treatment) to wane, the benefits may be offset if the contaminated quarantine pasture is subsequently used in the grazing rotation. Nevertheless, increasing the duration of quarantine, may be useful where the likelihood of establishment of AR populations of GINs is high and subsequent use of the quarantine area can be carefully managed. For example, bought-in cattle with high FECs and multiple anthelmintic treatment-resistant genotypes pose a challenge, particularly when pasture contamination by L3 is low on pastures grazed by the resident herd e.g. typically late winter/early spring. In such cases, the population of GINs *in refugia* are likely to be relatively small. These risks are heightened when purchasing from regions with suspected high levels of AR (although accurate estimates of regional prevalence are currently not available; Rose Vineer et al., 2022).

Taken together, our simulation results suggest that if quarantine duration is extended to reduce the risk of introducing AR to the resident herd, then cattle should be housed/yarded for quarantine, or the quarantine pasture should be removed from the grazing rotation until L3 contamination has waned to levels deemed acceptable to the farmer. Our simulations suggest a half-life of many months on pasture, often *>* 1 year, although this varies widely with season and geographic location. Although this half-life estimate is rather arbitrary for farmers in the absence of convenient methods for estimating pasture contamination, further integration of GLOWORMMETA with pasture contamination mapping frameworks, as suggested by McFarland et al.(2022), could provide farmers with useful data to evaluate competing biosecurity and grazing strategies to manage risk of introducing AR.

The model replicated population-level measures of intensity of infection in a range of groups of cattle well, demonstrating that continuous, mean-field models such as GLOWORM-META are useful for comparing candidate control strategies on *overall* infection levels. Care must be taken with the interpretation and application of such models, as they do not (and are not intended to) capture the range of intensities of infection observed within populations. For example, the mean FEC for cattle in the McFarland dataset (McFarland et al., 2022) was relatively low throughout the grazing season, but individual FECs of around 400 eggs per gram were observed. Models aiming to accurately replicate the distribution of infection within host populations (e.g. to inform targeted selective treatment, Greer et al. (2009); Berk et al. (2016)) would need to incorporate additional stochasticity to capture phenotypic plasticity Brass et al. (2021) and host factors (Vagenas et al., 2007). However, the benefits of this approach are offset against a potential lack of data to adequately parameterise the models (Verschave et al., 2014, 2016), incomplete understanding of the mechanisms driving the stochasticity (Berk et al., 2016), and computational requirements. For example, a similarly well-validated model was developed for *O. ostertagi* which incorporated parasite-induced anorexia, but included a simplified representation of the parasitic stages (Filipe et al., 2023). Furthermore, with increasing model complexity, effective continuous testing during model development becomes increasingly difficult, compromising rigour. Instead, GLOWORM-META incorporates both environmental stochasticity - through seasonally forced temperature - and rainfall as levels of precipitation. We used dependent variables, but opted to simulate average host characteristics, using a simple growth curve and dry matter intake estimates, for parsimony and computational reasons.

In conclusion, the aim of this current study was to develop a model framework, GLOWORM-META, which integrates the free-living and the parasitic stages of both *O. ostertagi* and *C. oncophora*, and host movements within a farm. Model simulations yielded insights into the relative influence of biosecurity practices on a range of outcomes of interest. These simulations suggest that biosecurity recommendations and further research intended to reduce the risk of introducing GIN populations (including AR populations) onto a farm could focus initially on practical recommendations for quarantine duration, safely reducing FECs during quarantine, and improving understanding of interactions with the season of purchase. Aspects of GIN biosecurity not evaluated in this study include the impact of quarantine treatments and existing pasture contamination (*refugia*) on selection for AR and explicit simulation of the potential introduction of resistant genotypes onto a farm. Models developed explicitly to track genotypes within sub-populations on pastures would allow this to be explored over a range of weather and climatic conditions, and over extended periods, which could then inform targeted empirical work to establish a stronger evidence base for practical recommendations. GLOWORM-META provides a basis for this further work, by establishing a well-validated, openly available framework for the research community.

## Appendix A. GLOWORM-META: Modelling gastrointestinal nematode metapopulation dynamics to inform cattle biosecurity research

**Figure A.13:**
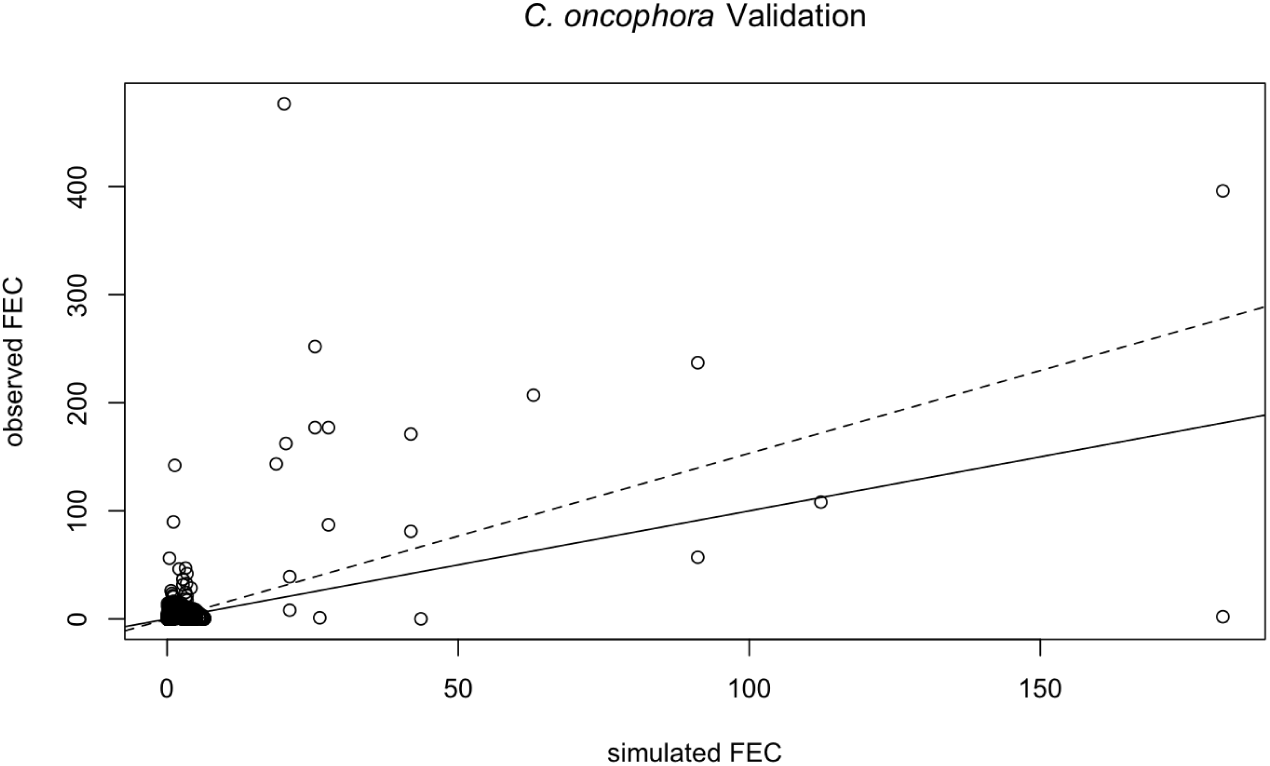
The observed and simulated faecal egg counts (points) for *C. oncophora* for the data used in the model validation. The solid black line indicates hypothetical perfect agreement between the observed and simulated faecal egg counts. The grey dashed line indicates the predicted slope of the regression.

**Figure A.14:**
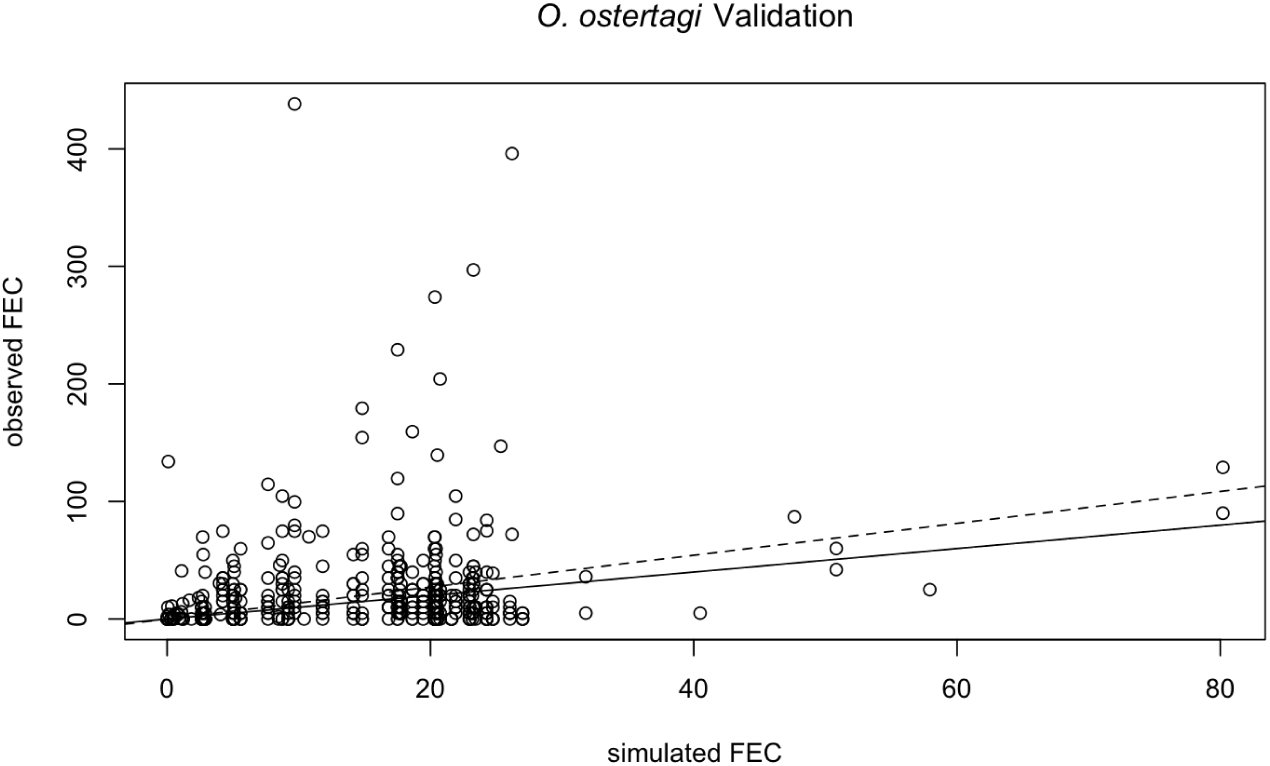
The observed and simulated faecal egg counts (points) for *O. ostertagi* for the data used in the model validation. The solid black line indicates hypothetical perfect agreement between the observed and simulated faecal egg counts. The grey dashed line indicates the predicted slope of the regression.

**Figure A.15:**
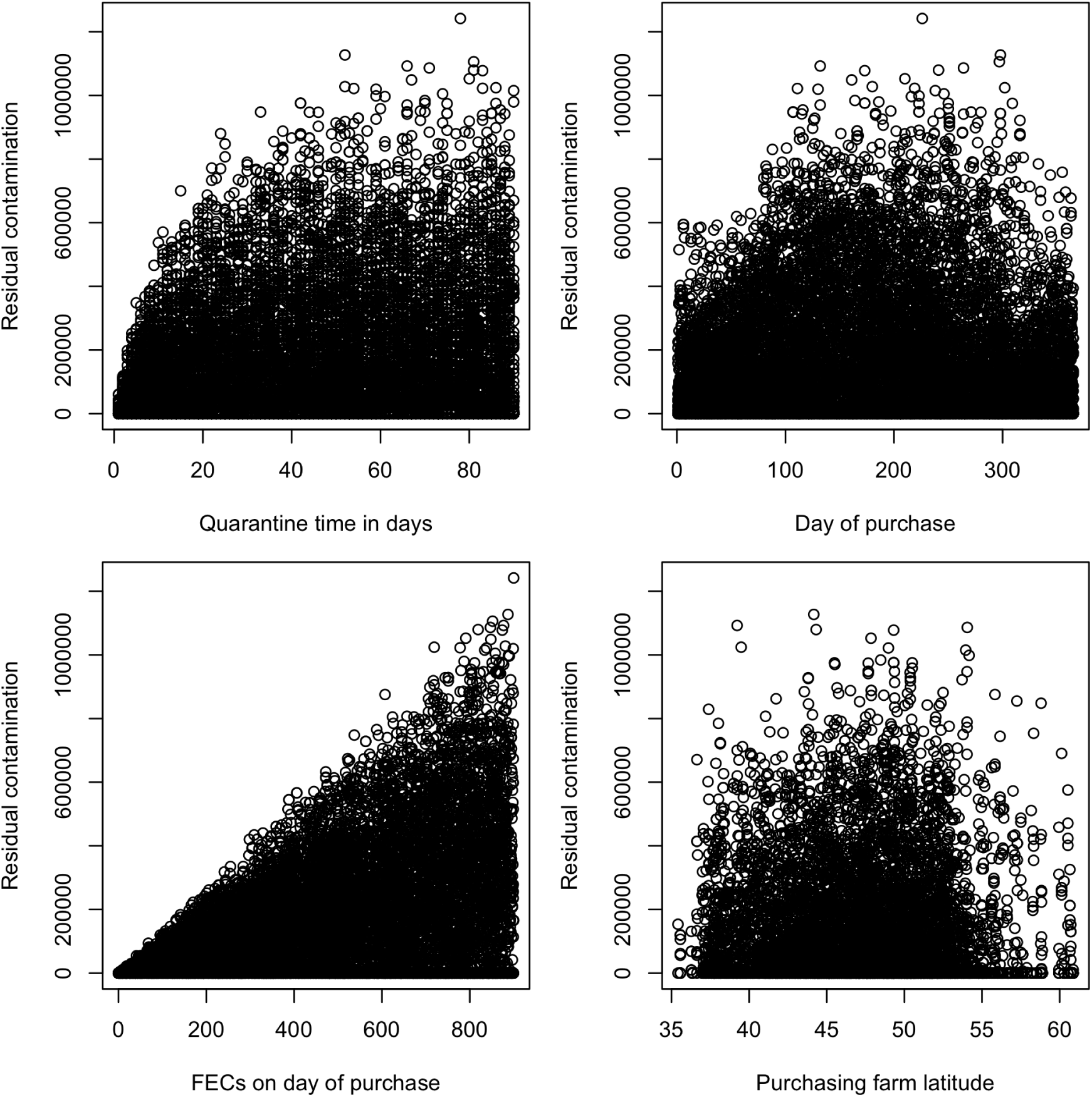
Distribution of residual contamination for *C. oncophora* simulations in Europe against the four parameters used in each of the simulations.

**Figure A.16:**
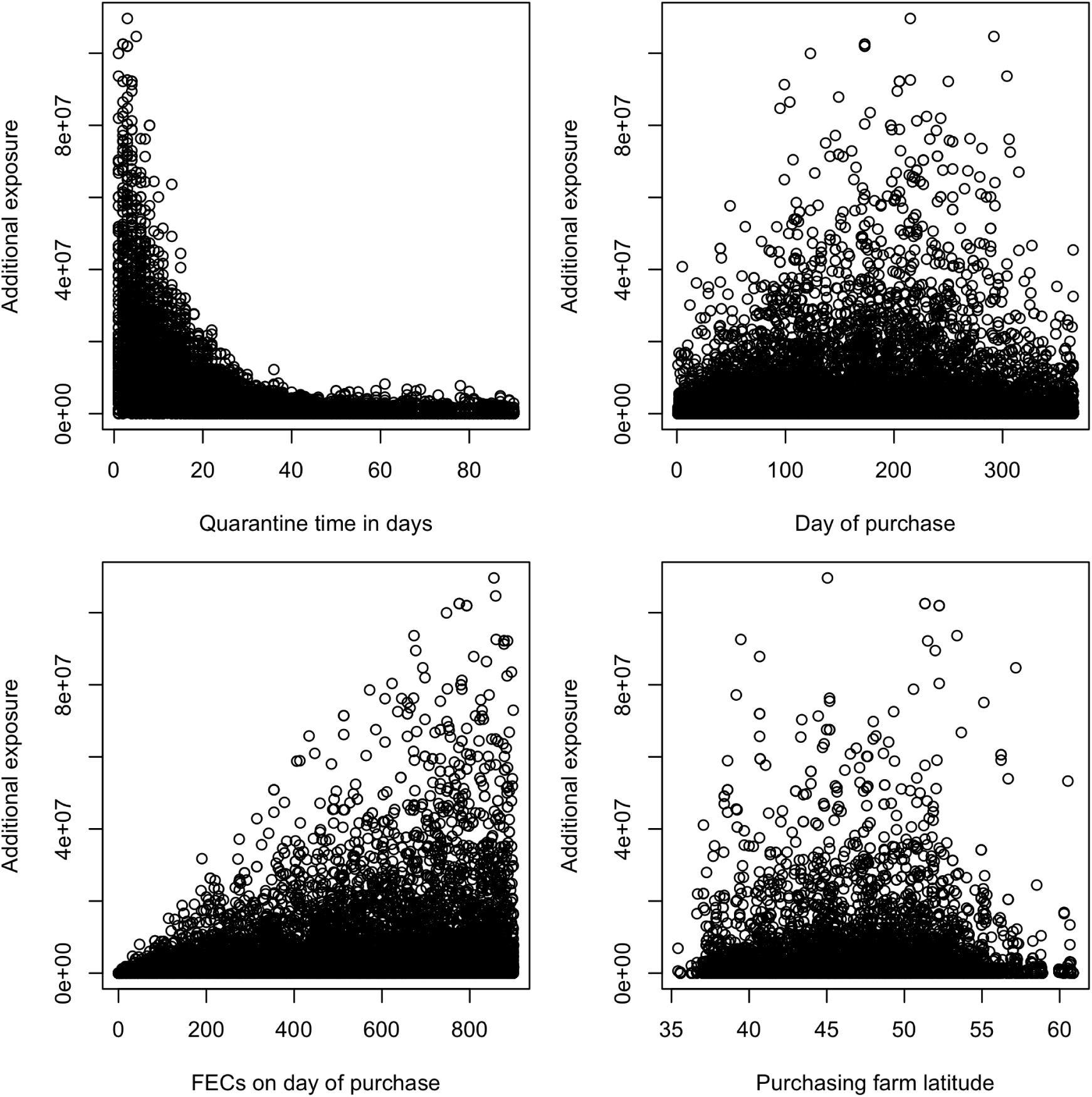
Distribution of additional exposure for *C. oncophora* simulations in Europe against the four parameters used in each simulation

**Figure A.17:**
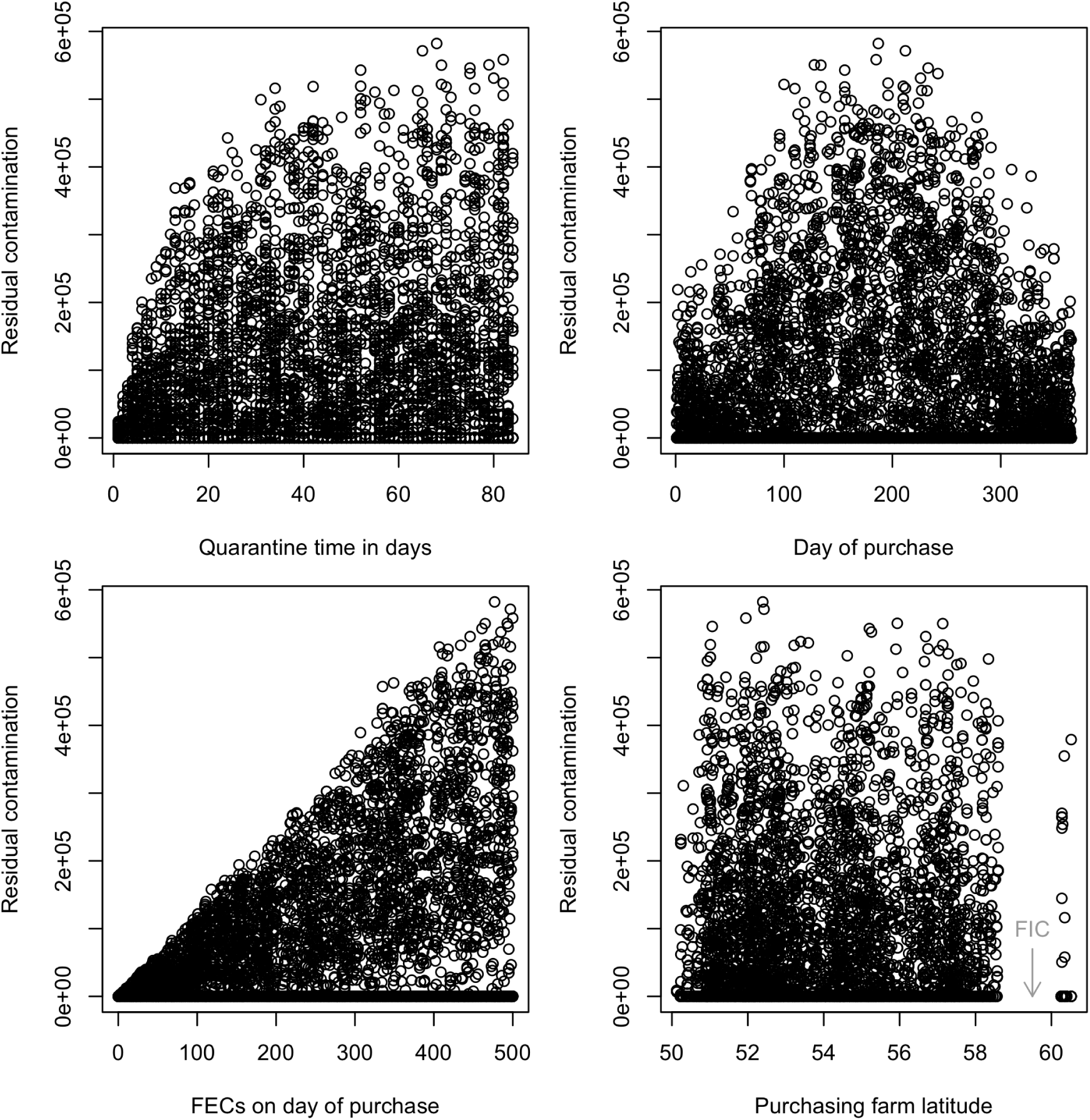
Distribution of residual contamination for *C. oncophora* simulations in the UK against the four parameters

**Figure A.18:**
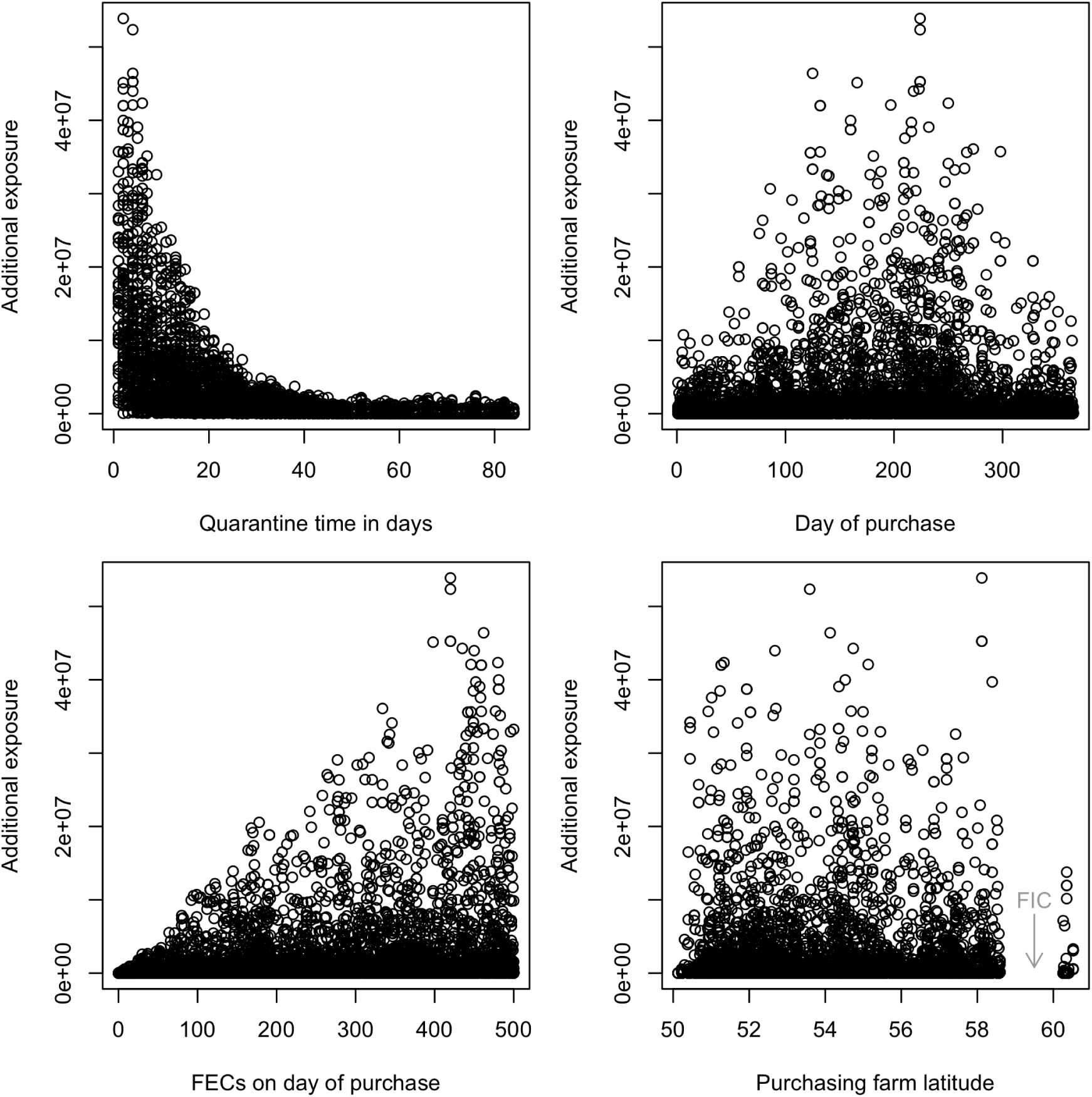
Distribution of additional exposure for *C. oncophora* simulations in the UK against the four parameters

**Figure A.19:**
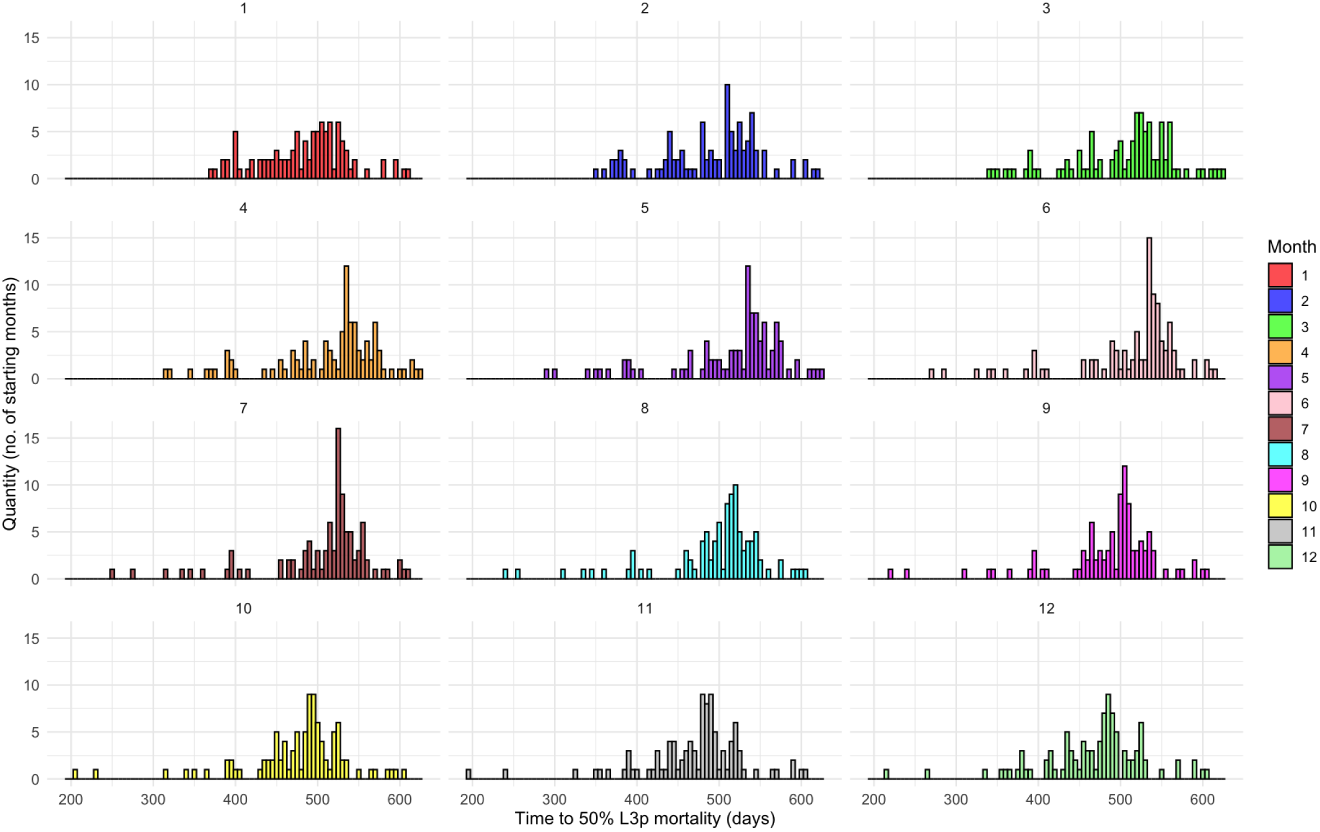
Distribution of the number of the months against larval mortality duration for *O. ostertagi*. Months are coloured as per the key.

**Figure A.20:**
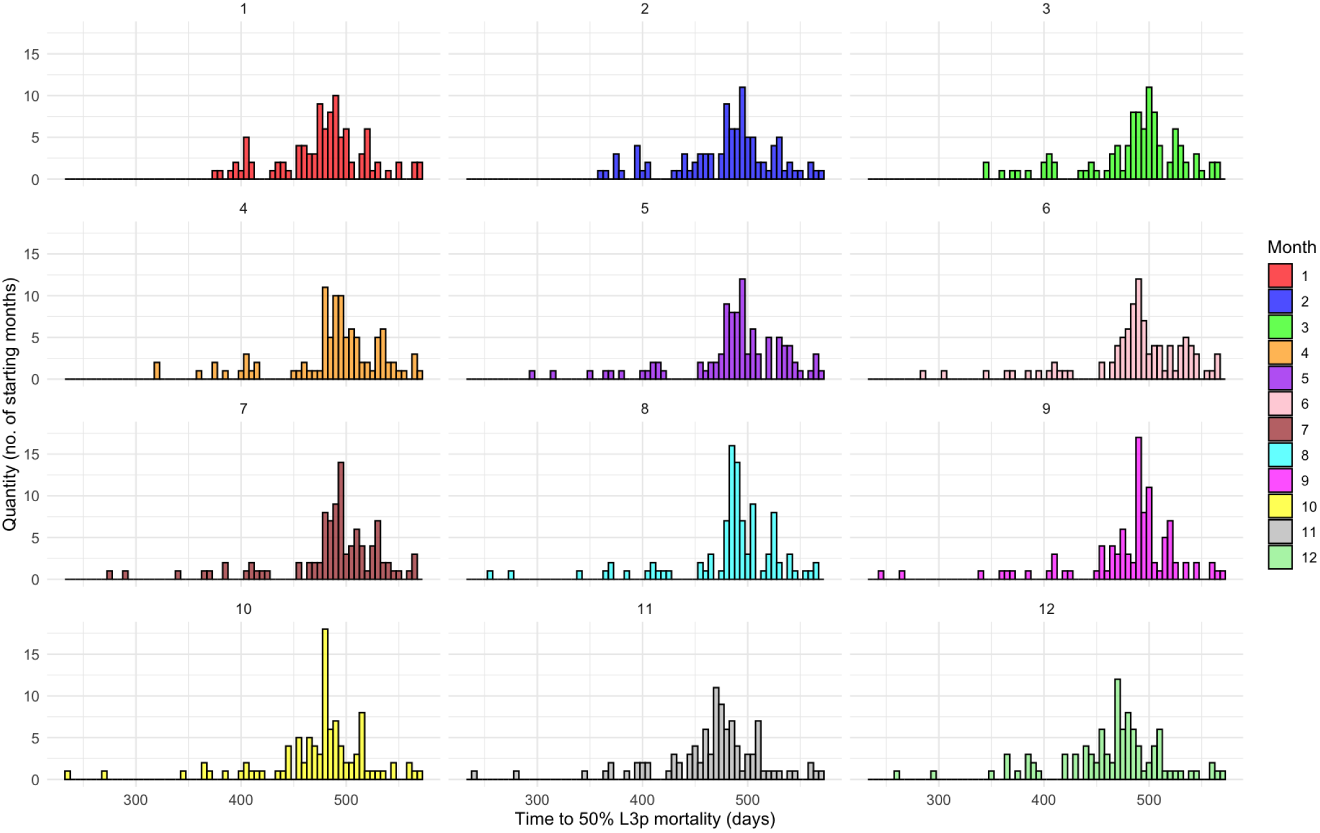
Distribution of the number of the months against larval mortality duration for *C. oncophora*. Months are coloured as per the key.

**Table A.2:** Summary of the validation data used for GLOWORM-META.

| Study | Herd | Location | Number of animals | Age of animals (months) | Start Date | End Date | Stocking Rate | Anthelmintic Used? | Treatment Date | Anthelmintic Used | Grass Quality (kgDM) |
| --- | --- | --- | --- | --- | --- | --- | --- | --- | --- | --- | --- |
| Verschave et al, 2015 | 1 | Dudzele, BE | 11 | 20 | 20/04/2012 | 09/11/2012 | 5.2 | Yes | 07/09/2012 | Moxidectin | 2000 |
|  | 2 | Malle, BE | 12 | 6 | 15/05/2012 | 06/09/2012 | 10 | Yes | 07/09/2012 | Doramectin | 2000 |
|  | 3 | Evergem, BE | 10 | 21 | 15/05/2012 | 06/09/2012 | 5.2 | No | – | – | 2000 |
|  | 4 | Oudenaarde, BE | 19 | 19 | 20/04/2013 | 26/11/2013 | 13 | No | – | – | 2000 |
|  | 5 | Drongen, BE | 11 | 11 | 07/06/2013 | 14/10/2013 | 62.4 | No | – | – | 2000 |
|  | 6 | Sinaai, BE | 16 | 15 | 20/06/2013 | 30/11/2013 | 3.7 | No | – | – | 2000 |
|  | 7 | Eeklo, BE | 16 | 10 | 13/06/2013 | 10/09/2013 | 25.5 | Yes | 19/08/2013 | Moxidectin | 2000 |
| Passage Herd | Co | Penciuk, GB | 2 | 12 | 24/05/2023 | 30/09/2023 | 9 | No | – | – | 3054 |
|  | Oo | Penciuk, GB | 2 | 12 | 24/05/2023 | 30/09/2023 | 8.84 | No | – | – | 3826 |
| McFarland et al, 2022 | A | Hillsborough, NI | 11 | 12 | 19/04/2019 | 19/04/2020 | 40 | Yes | 28/05/2019 | Doramectin | 3000 |
|  | B | Hillsborough, NI | 11 | 12 | 19/04/2019 | 19/04/2020 | 40 | Yes | 28/05/2019 | Doramectin | 3000 |
|  | C | Hillsborough, NI | 11 | 12 | 19/04/2019 | 19/04/2020 | 40 | Yes | 28/05/2019 | Doramectin | 3000 |
|  | D | Hillsborough, NI | 11 | 12 | 19/04/2019 | 19/04/2020 | 40 | Yes | 28/05/2019 | Doramectin | 3000 |

## References

Adler, D., Nenadic, T., Kotenko, A., 2023. vioplot: Violin Plot. URL: https://CRAN.R-project.org/package=vioplot. r package version 0.4.0. for Animal Health, W.O., 2019. Terrestrial Animal Health Code: Guidelines on the Prevention and Control of Parasitic Diseases in Livestock.

Barnes, E., Dobson, R., 1990. Population dynamics of trichostrongylus colubriformis in sheep: Computer model to simulate grazing systems and the evolution of anthelmintic resistance. International Journal of Parasitology 20, 823 – 831.

Bates, A.J., Greer, A., McAnulty, R., Jackson, A., 2012. Targeted selective treatment with anthelmintic for new zealand dairy heifers. Veterinary Parasitology 309.

Berk, Z., Bishop, S.C., Forbes, A.B., Kyriazakis, I., 2016. A simulation model to investigate interactions between first season grazing calves and ostertagia ostertagi. Veterinary Parasitology 226, 198 – 209.

Bolajoko, M.B., Rose, H., Musella, V., Bosco, A., Rinaldi, L., van Dijk, J., Cringoli, G., Morgan, E.R., 2015. The basic reproduction quotient (q0) as a potential spatial predictor of the seasonality of ovine haemonchosis. Hydrological Processes 9, 333 – 350.

Brass, D.P., Cobbold, C.A., Ewing, D.A., Purse, B.V., Callaghan, A., White, S.M., 2021. Phenotypic plasticity as a cause and consequence of population dynamics. Ecology Letters 24.

Bricarello, P.A., da Rocha, R.A., Longo, C., Hötzel, M.J., 2023. Understanding animal-plant-parasite interactions to improve the management of gastrointestinal nematodes in grazing ruminants. Pathogens 12, 531.

Charlier, J., Höglund, J., Morgan, E.R., Geldhof, P., Vercruysse, J., Claerebout., E., 2020. Biology and epidemiology of gastrointestinal nematodes in cattle. Veterinary Clinics: Food Animal Practice 36, 1 – 15.

Coop, R.L., Kyriazakis, I., 2001. Influence of host nutrition on the development and consequences of nematode parasitism in ruminants. Trends in Parasitology 17, 325–330.

COWS, 2021. Best Practice Guidelines for Worm Control in Cattle.

Damiaans, B., Sarrazin, S., Heremans, E., Dewulf, J., 2018. Perception, motivators and obstacles of biosecurity in cattle production. Vlaams Diergeneeskundig Tijdschrift 87, 150–163.

van Dijk, J., David, G, P., Baird, G., Morgan, E, R., 2008. Back to the future: Developing hypotheses on the effects of climate change on ovine parasitic gastroenteritis from historical data. Veterinary Parasitology 158, 73 – 84.

Dobson, R., Barnes, E., Tyrrell, K., Hosking, B., Larsen, J., Besier, R., Love, S., Rolfe, P., Bailey, J., 2011. A multi-species model to assess the effect of refugia on wormcontrol and anthelmintic resistance in sheep grazing systems. Australian Veterinary Journal 89, 200 – 208.

Doe, J., Smith, A., Lee, B., 2024. DataScienceR: Tools for Data Science in R. URL: https://CRAN.R-project.org/package=DataScienceR. r package version 1.2.3.

E-OBS, 2023. E-OBS: Daily gridded meteorological data for Europe, Version 29. European Climate Assessment & Dataset (ECA&D). URL: https://www.ecad.eu/download/ensembles/.

Filipe, J., Kyriazakis, I., McFarland, C., Morgan, E., 2023. Novel epidemiological model of gastrointestinal nematode infection to assess grazing cattle resilience by integrating host growth, parasite, grass and environmental dynamics. International Journal for Parasitology 53, 133 – 155.

Gethings, O., Rose, H., Mitchell, S., Van Dijk, J., Morgan, E., 2015. Asynchrony in host and parasite phenology may decrease disease risk in livestock under climate warming: Nematodirus battus in lambs as a case study. Parasitology 142, 1306 – 1317.

Greer, A., Kenyon, F., Bartley, D., Jackson, E., Gordon, Y., Donnan, A., McBean, D., Jackson, F., 2009. Development and field evaluation of a decision support model for anthelmintic treatments as part of a targeted selective treatment (tst) regime in lambs. Veterinary Parasitology 164, 12 – 20.

Hijmans, R.J., 2024. geosphere: Spherical Trigonometry. URL: https://github.com/rspatial/geosphere. r package version 1.5-20.

Hoste, H., Torres-Acosta, J., 2011. Non chemical control of helminths in ruminants: Adapting solutions for changing worms in a changing world. Veterinary Parasitology 180, 144–154.

Iooss, B., Weber, F., 2016. soboljansen: Monte Carlo Estimation of Sobol’ Indices. URL: https://cran.r-project.org/web/packages/soboljansen/index.html. r package version 1.0.0.

Jackson, R., Rhodes, A., Pomroy, W., Leathwick, D., West, D., Waghorn, T., Moffat, J., 2011. Anthelmintic resistance and management of nematode parasites on beef cattle-rearing farms in the north island of new zealand. Veterinary Clinics of North America: Food Animal Practice 54, 289 – 296.

Kaplan, R.M., 2020. Biology, epidemiology, diagnosis, and management of anthelmintic resistance in gastrointestinal nematodes of livestock. Veterinary Clinics of North America: Food Animal Practice 36, 17 – 30.

Kelleher, A.C., Good, B., de Waal, T., Keane, O.M., 2020. Anthelmintic resistance among gastrointestinal nematodes of cattle on dairy calf to beef farms in ireland. Irish Veterinary Journal 73.

Khanyari, M., Oyanedel, R., Khara, A., Sharma, M., E J Milner-Gulland, K.R.S., Vineer, H.R., Morgan, E.R., 2024. Predicting and reducing potential parasite infection between migratory livestock and resident asiatic ibex of pin valley, india. Springer: Journal of Biosciences 49.

Lancaster, M., 1970. The recovery of infective nematode larvae from herbage samples. Journal of Helminthology 44, 219 – 230.

Lubinda, J., Yaxin Hamainza, B., Haque, U., Moore, A.J., 2021. Modelling of malaria risk, rates, and trends: A spatiotemporal approach for identifying and targeting sub-national areas of high and low burden. PLOS Computational Biology, 1 – 23.

Martínez-Valladares, M., Geurden, T., Bartram, D., Martínez-Pérez, J., Robles-Pérez, D., Bohórquez, A., Florez, E., Meana, A., Rojo-Vázquez, F., 2015. Resistance of gastrointestinal nematodes to the most commonly used anthelmintics in sheep, cattle and horses in spain. Veterinary Parasitology 211.

Mayo, C., McDermott, E., Kopanke, J., Stenglein, M., Lee, J., Mathiason, C., Carpenter, M., Reed, K., Perkins, T.A., 2020. Ecological dynamics impacting bluetongue virus transmission in north america. Frontiers 7.

McCarthy, C., Vineer, H.R., Morgan, E.R., van Dijk, J., 2022. Predicting the unpredictable? a climate-based model of the timing of peak pasture infectivity for dictyocaulus viviparus. Veterinary Parasitology 309. 109770.

McFarland, C., Vineer, H.R., Chesney, L., Henry, N., Brown, C., Airs, P., Nicholson, C., Scollan, N., Lively, F., Kyriazakis, I., Morgan, E.R., 2022. Tracking gastrointestinal nematode risk on cattle farms through pasture contamination mapping. International Journal of Parasitology 52, 691–703.

McGowan, T.A., O’Leary, M.K., 2021. naturalearth: Natural Earth Data for R. URL: https://cran.r-project.org/web/packages/naturalearth/index.html. r package version 0.1.0.

Milewski, S., Gümus, T., 2023. Nutritional status and immune response interactions in grazing livestock. Animals 13, 731.

Papadopoulos, E., Gallidis, E., Ptochos, S., 2012. Anthelmintic resistance in sheep in europe: A selected review. Veterinary Clinics of North America: Food Animal Practice 189, 85 – 88.

Pebesma, E., Bivand, L.D., van D. M. R.B.R.A.J., 2023. sf: Simple Features for R. URL: https://cran.r-project.org/web/packages/sf/index.html. r package version 1.0-0.

Pierce, D., 2019. ncdf4: Interface to Unidata netCDF (Version 4 or Earlier) Format Files. R package version 1.22, Available at https://CRAN.R-project.org/package=ncdf4.

Ploeger, H, W., Kloosterman, A., Rietveld, F, W., 1995. Acquired immunity against cooperia spp. and ostertagia spp. in calves: effect of level of exposure and timing of the midsummer increase. Veterinary Parasitology 58, 61 – 74.

R Core Team, 2023. R: A Language and Environment for Statistical Computing. R Foundation for Statistical Computing. Vienna, Austria. Version 4.3.0, Available at https://www.R-project.org/.

Rose, H., Wang, T., van Dijk, J., Morgan, E.R., 2015. Gloworm-fl: A simulation model of the effects of climate and climate change on the free-living stages of gastro-intestinal nematode parasites of ruminants. Ecological Modelling 197, 232 – 245.

Rose, J., 1961. Some observations on the free-living stages of ostertagia ostertagi, a stomach worm of cattle. Parasitology 51, 295–307.

Rose Vineer, H., Morgan, E.R., Hertzberg, H., Bartley, D.J., Bosco, A., Charlier, J., Chartier, C., Claerebout, E., de Waal, T., Hendrick, G., Hinney, B., H€oglund, J., Jězek J., Kasny, M., Keane, O.M., Martınez-Valladares, M., Mateus, T.L., McIntyre, J., Mickiewicz, M., Munoz, A.M., Phythian, C.J., Ploeger, H.W., Rataj, A.V., Skuce, P.J., Simin, S., Sotiraki, S., Spinu, M., Stuen, S., Thamsborg, S.M., Vadlejch, J., Varady, M., von Samson-Himmelstjerna, G., Rinaldi, L., 2020a. Increasing importance of anthelmintic resistance in european livestock: creation and meta-analysis of an open database. Parasite 27.

Rose Vineer, H., Verschave, S., Claerebout, E., Vercruysse, J., Shaw, D., Charlier, J., Morgan, E., 2020b. Gloworm-para: a flexible framework to simulate the population dynamics of the parasitic phase of gastrointestinal nematodes infecting grazing livestock. International Journal of Parasitology 5, 883–885.

Sauermann, C.W., Leathwick, D.M., 2018. A climate-driven model for the dynamics of the free-living stages of cooperia oncophora. Veterinary Parasitology 255, 83 – 90.

Smith, G., Grenfell, B.T., Isham, V., Cornell, S., 1999. Anthelmintic resistance revisited: under-dosing, chemoprophylactic strategies, and mating probabilities. International Journal of Parasitology 29, 77 – 91.

Soetaert, K., Petzoldt, T., Setzer, R.W., 2010. deSolve: Solvers for Initial Value Problems of Differential Equations. R package version 1.34, Available at https://CRAN.R-project.org/package=deSolve.

Szmaragd, C., Wilson, A.J., Carpenter, S., Wood, J.L.N., Mellor, P.S., Gubbins, S., 2009. A modeling framework to describe the transmission of bluetongue virus within and between farms in great britain. Frontiers 4.

Taylor, M., 2012. Scops and cows—‘worming it out of uk farmers’. Veterinary Parasitology 186, 65 – 69.

Vagenas, D., Bishop, S.C., Kyriazakis, I., 2007. A model to account for the consequences of host nutrition on the outcome of gastrointestinal parasitism in sheep: logic and concepts. Parasitology 134, 1263 – 1277.

Vercruysse, J., Claerebout, E., 1997. Immunity development against ostertagia ostertagi and other gastrointestinal nematodes in cattle. Veterinary Parasitology 72, 309–326.

Verschave, S., 2015. Development of a transmission model for gastro-intestinal nematode infections in cattle. Ph.D. thesis. Ghent University.

Verschave, S., Vercruysse, J., Claerebout, E., Rose, H., Morgan, E., Charlier, J., 2014. The parasitic phase of ostertagia ostertagi: quantification of the main life history traits through systematic review and meta-analysis. International Journal for Parasitology 44, 1091 – 1104.

Verschave, S.H., Rose, H., Morgan, E.R., Claerebout, E., Vercruysse, J., Charlier, J., 2016. Modelling cooperia oncophora: Quantification of key parameters in the parasitic phase. Veterinary Parasitology 223, 111 – 114.

Vineer, H.R., 2020. What modeling parasites, transmission, and resistance can teach us. Veterinary Clinics: Food Animal Practices 36, 145 – 158.

Walker, J.G., Evans, K.E., Vineer, H.R., van Wyk, J.A., Morgan, E.R., 2018. Prediction and attenuation of seasonal spillover of parasitesbetween wild and domestic ungulates in an arid mixed-use system. Journal of Applied Ecology 55, 1976–1986.

Wang, T., Avramenko, R.W., Redman, E.M., Wit, J., Gilleard, J.S., Colwell, D.D., 2020. High levels of third-stage larvae (l3) overwinter survival for multiple cattle gastrointestinal nematode species on western canadian pastures as revealed by its2 rdna metabarcoding. Parasites & Vectors 13, 458.

Wang, T., Vineer, H.R., Redman, E., Morosetti, A., Chen, R., McFarland, C., Colwell, D.D., Morgan, E.R., Gilleard, J.S., 2022. An improved model for the population dynamics of cattle gastrointestinal nematodes on pasture: parameterisation and field validation for ostertagia ostertagi and cooperia oncophora in northern temperate zones. Veterinary Parasitology 310.

Wickham, H., 2011. testthat: Get started with testing. The R Journal 3, 5 – 10.

Wood, S.N., 2017. mgcv: Generalized Additive Models. URL: https://cran.r-project.org/web/packages/mgcv/index.html. accessed: 2025-02-05.

Xu, C., Singh, V.P., 2001. Evaluation and generalization of temperature-based methods for calculating evaporation. Hydrological Processes 15, 305 – 319.

